# Decoding genetic architecture of dog complex traits by constructing fine-scale genomic ancestry of admixture

**DOI:** 10.1101/2023.09.17.558101

**Authors:** Shilei Zhao, Guo-Dong Wang, Yanhu Liu, Ya-Ping Zhang, Hua Chen

**Affiliations:** Beijing Institute of Genomics, Chinese Academy of Sciences and China National Center for Bioinformation, Beijing 100101, China; State Key Laboratory of Genetic Resources and Evolution, Kunming Natural History Museum of Zoology, Kunming Institute of Zoology, Chinese Academy of Sciences, Kunming 650201, China; University of Chinese Academy of Sciences, Beijing 100049, China; Center for Excellence in Animal Evolution and Genetics, Chinese Academy of Sciences, Kunming 650223, China

**Author notes:** Correspondence: Hua Chen. These authors contributed equally (Shilei Zhao, Guo-Dong Wang).

**Keywords:** admixture, genomic ancestry, genetic architecture, complex traits, recombination, hidden Markov model

## Abstract

Domestic animals and plants exhibit remarkable phenotypic diversity in terms of morphology, behavior, and physiology, which can be attributed to the complex interbreeding process of various breeds and artificial selection. Here we develop a method that can efficiently construct fine-scale interbreeding history of local segments along the genome. Since ancestral breeds usually exhibit diverse phenotypes, the method provides a valuable approach for unraveling the genetic architecture of complex traits in admixed breeds. Simulated data demonstrates that the method performs well, even in scenarios involving complex interbreeding with up to 19 ancestral breeds.

The method is applied to analyze three mixed dog breeds, Irish Wolfhound, Giant Schnauzer, and Miniature Schnauzer, representing instances of body-size enlargement and miniaturization. Numerous novel ancestor breed-specific genes determining body size are identified, including *FGFR2, WDR11*, and *FARS2*. We also validate genes reported in previous GWAS or genomic sweep scans, such as *LCORL, STC2, NPR2*, and *FGF4*. These findings highlight the validity of the method as a valuable tool for investigating the genetic basis underlying ancestry-specific traits in domestic animals and plants with complex interbreeding histories.

## Introduction

Modern domesticated dogs have been sculpted by artificial selection for at least 14,000 years ^1, 2, 3, 4^. Two distinct artificial selection mechanisms have played a crucial role in shaping the genetic diversity of domesticated dogs: intense artificial selection on newly occurring mutations led to the development of novel phenotypes; in addition, the interbreeding of different breeds transferred existing mutations from a breed with unique phenotypes to another breed with different genetic background, rapidly generating a high degree of phenotype diversity ^5^. The breed formation process was accelerated during the Victorian era (circa 1830-1900) through the implementation of systematic breeding practices, especially with the establishment of breed clubs. The intensive breeding programs during that period produced more than 400 modern breeds, showcasing unrivaled diversity in morphological, behavioral, and physiological phenotypes. The remarkable phenotypic diversity in domestic dogs, coupled with their distinctive interbreeding process, provides an inimitable system for studying the genetics of mammalian traits ^6^ and furthermore, provide insights into human health and phenotype as well ^7^.

Extensive genome-wide association studies (GWAS) and selective sweep scans have successfully identified multiple functional variants associated with morphologic traits, disease susceptibility, and behavior ^8, 9, 10^, such as coat type ^11^, body size ^10, 12, 13, 14^, ear floppiness ^9^, skin wrinkling ^15^, lactase persistence ^16^, and antiparasite ability ^17^. A well-known example is *IGF1*, identified through a selective sweep scan, which determines body size miniaturization among small dog breeds ^12^.

However, none of these studies utilized the complex interbreeding history from the artificial selection and admixture perspective. Reconstructing interbreeding history can be informative for understanding the origin and mechanism underlying breed-specific phenotypes. In addition, it is common that ancestral breeds are shared among multiple admixed dog breeds, allowing for a replicated experimental design with varying interbreeding history and linkage disequilibrium (LD) levels. Leveraging the shared ancestral breeds from different admixed breeds can mitigate false positives caused by breeding bottlenecks and enables identification of functional regions in a fine scale.

Human genetic studies have successfully applied admixture mapping approach to identify genes associated with ancestry-specific diseases in recently admixed populations ^18^. However, there are challenges in applying this approach to mixed-breed dogs. First, existing methods for inferring local ancestry segments are primarily designed for admixed populations with a limited number of ancestors ^19, 20^, which may be less effective for dogs and other domestic animals with complex and long-time admixture histories. Second, intensive artificial selection and strict breeding systems in dogs have led to reduced intra-breed phenotypic and genotypic heterogeneity, making current individual-based admixture mapping methods less suitable for dogs.

In this paper, we present a new method named GAMA (Genomic Analysis of Multiple Admixture) for accurately decoding the fine-scale ancestral origins of chromosome segments in the genome of admixed animal breeds or plant lines. We demonstrate with simulation that GAMA outperforms existing methods, particularly in scenarios involving multiple ancestral populations and/or ancient admixture events. Additionally, GAMA is computationally efficient and scalable to thousands of samples. We apply GAMA to three cases of admixed dog breeds, including Irish Wolfhound, Giant Schnauzer, and Miniature Schnauzer. These breeds were developed to change body sizes by transfer existing mutations controlling body size from different parental breeds to a baseboard breed. Irish Wolfhound and Giant Schnauzer share one ancestral population, namely the Great Dane. Giant Schnauzers and Miniature Schnauzers have the same baseboard ancestor, Standard Schnauzer, but with opposite breeding directions of body size (enlarging and miniaturizing). Our analysis identifies numerous novel ancestral breed-specific genes contributing to body size, including *FGFR2, WDR11* and *FARS2* contributing to enlargement, and *FGF4, FGF5, TFAP2B, BCL6, FGFR3* contributing to miniaturization. Interestingly, a significant overlap of enlargement genes with the Great Dane origin is identified in Irish Wolfhound and Giant Schnauzer. Overall, this study presents a novel approach for constructing fine-scale genomic ancestry and investigating the genetic basis of complex traits in admixed breeds or populations. The proposed method serves as a valuable complement to the existing association study and selective sweep methods, and enhances our understanding of genetic architecture of complex traits in domestic animals and plants.

## Results

### Performance of inferring local ancestors and parameters

We simulate genomic data from 3-way, 5-way, and 7-way admixture mimicking the possible interbreeding history of Giant Schnauzer, Irish Wolfhound and Miniature Schnauzer. We also simulate a 19-way admixture to evaluate the performance of GAMA in the cases of complex admixture with numerous ancestral populations (see details of each scenario in supplementary information). In addition to using natural ancestral populations as reference populations, we also use wolf as a reference population to test the robustness of GAMA for inferring local ancestry segments when some ancestral populations are unknown, which are common in dog breeds.

The simulation data consist of sixty scenarios involving the combination of different admixture time (5, 10, 20, 50, and 100 generations) and locus-specific error rates (0, 0.02, and 0.05). Each simulated dataset includes 20 haplotypes sampled from the admixed population. We compare the performance of GAMA with the newly developed and widely used method Mosaic ^21^. Both GAMA and Mosaic are applied to the simulated data, and GAMA consistently outperform Mosaic across all parameter settings. The difference is particularly pronounced for long-term admixture time and large numbers of ancestral populations (Fig. 1). For instance, at the admixture time of λ=50 generations with no error rate (*α* = 0), the accuracies of GAMA are 0.9302, 0.9169, 0.8716, and 0.7841 for 3-way, 5-way, 7-way, and 19-way admixture respectively. In contrast, the accuracies of Mosaic are lower, measuring at 0.2596, 0.3593, 0.3619, and 0.2757 respectively (Fig. 1a). Both methods are robust against high error rates. Furthermore, GAMA is robust to using wolf as a reference ancestor, with a mean estimated ancestral proportion of the wolf being 1.9719e-04 across all simulations.

**Fig. 1.**
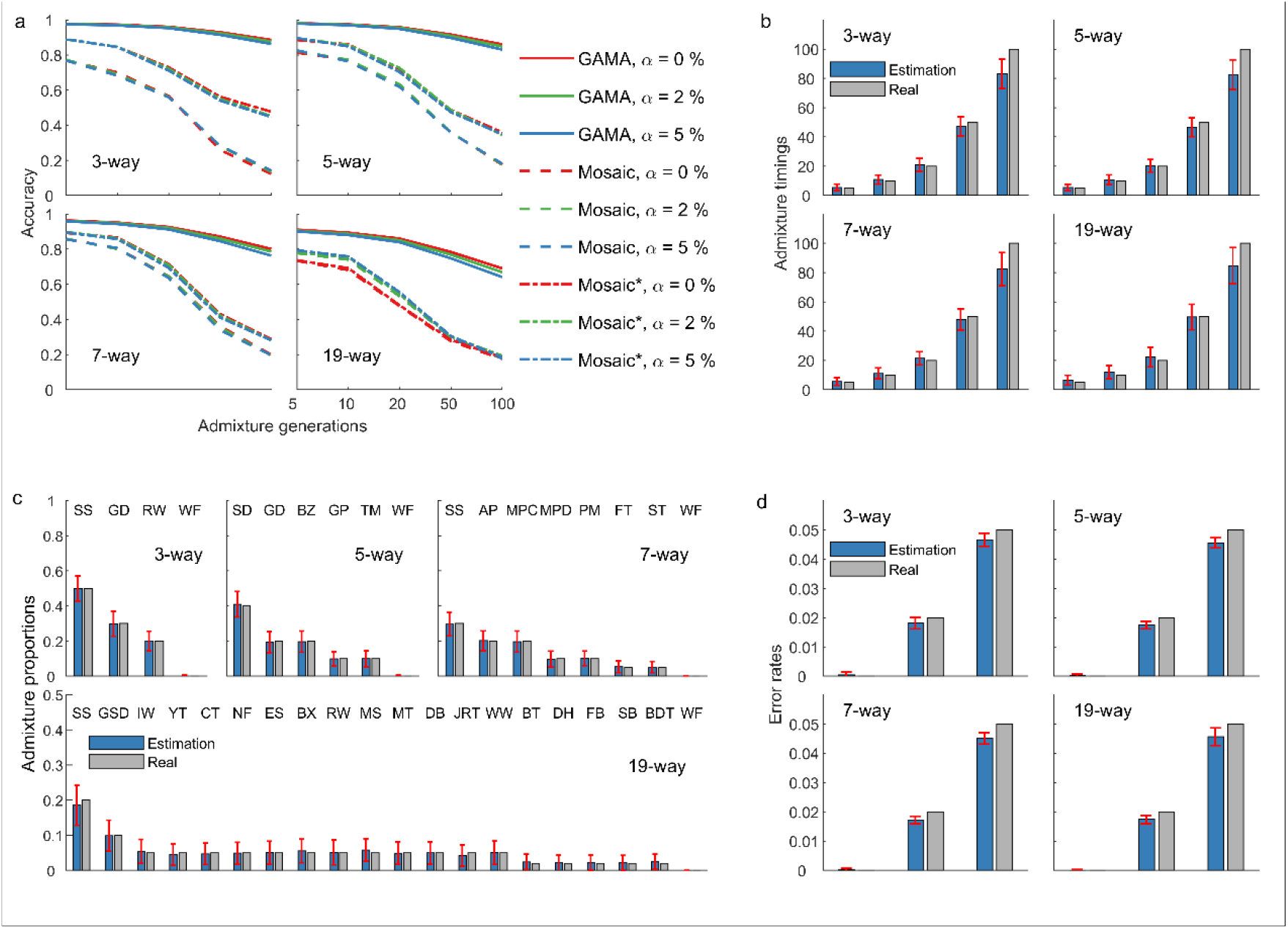
Performance of GAMA on interring local ancestral segments. **a** Accuracy of GAMA and Mosaic on inferring local ancestor on simulated data with different parameter settings including ancestral population number, admixture generation, and error rate. Four breeding schemes, including 3-way, 5-way, 7-way and 19-way admixtures were simulated. The details of ancestral populations are listed in Supplementary Table 1. Admixture time was set to be λ=5, 10, 20, 50, 100 generations, and error rates α=0, 0.02, 0.05. When running the two software, we set wolf as a pseudo ancestor and removed the results from Mosaic with pseudo ancestor are denoted as Mosaic*. **b**, Accuracy of GAMA on estimating admixture times. **c** Accuracy of GAMA on estimating admixture proportions. The abbreviated breed names are shown on top of the figures, and the full names are listed in Supplementary Table 1. **d** Accuracy of GAMA on estimating error rates. Red error bars represent -2 to 2 standard deviations from mean values.

However, the performance of Mosaic is affected by including wolf data in the analysis. After removing wolf as one of the reference ancestors, the performance of Mosaic in estimating the local ancestor is significantly boosted, with the accuracies of 0.5630, 0.4872, 0.4313, and 0.2837 for 3-way, 5-way, 7-way, and 19-way admixture respectively (error rate =0, Fig. 1a). GAMA can accurately infer parameters, including admixture times, error rates, and ancestor proportions (Fig. 1b-d). The estimation of error rates and ancestor proportions are unbiased. The estimations of admixture times are unbiased for short-term admixture times, while it tends to underestimate long-term admixture times (100 generations, Fig. 1b). The outliers in the estimations are mainly because of the small size of the simulated genome data (∼122Mb with 6,611 SNPs).

GAMA is computationally much more efficient than the existing methods. The mean running time for GAMA is 28.32, 36.6, 51.04, and 138.12 seconds for 3-way, 5-way, 7-way, and 19-way admixture, respectively (single thread mode, Intel(R) Xeon(R) Gold 5118 CPU @ 2.30GHz), and the mean running time for Mosaic is 164.43, 175.52, 279.71, and 2768.8 seconds respectively (with 20 threads).

Overall, GAMA performs well in inferring local genomic ancestries (especially for long-term admixture and multiple-way admixture) and parameters. The running time is acceptable for whole-genome and large-scale admixture populations. Thus, GAMA is suitable for studying domestic animals and plants with complex interbreeding histories.

### Constructing fine-scale genomic ancestry of three dog breeds

We apply GAMA to analyze three dog breeds developed by interbreeding of multiple ancestral breeds, including Irish Wolfhound, Giant Schnauzer, and Miniature Schnauzer. The three admixture cases all aimed to dramatically modify the body size of baseboard breeds while retaining their morphological characteristics such as coat type, ear morphology, and head morphology. The interbreeding history of the three breeds were documented clearly (Fig. 2, see Methods for details). Irish Wolfhound and Giant Schnauzer shared the same ancestral breed, Great Dane. Giant Schnauzer and Miniature Schnauzer originated from the same baseboard breed, Standard Schnauzer, with the contrary breeding direction of enlarging and reducing body size.

**Fig. 2.**
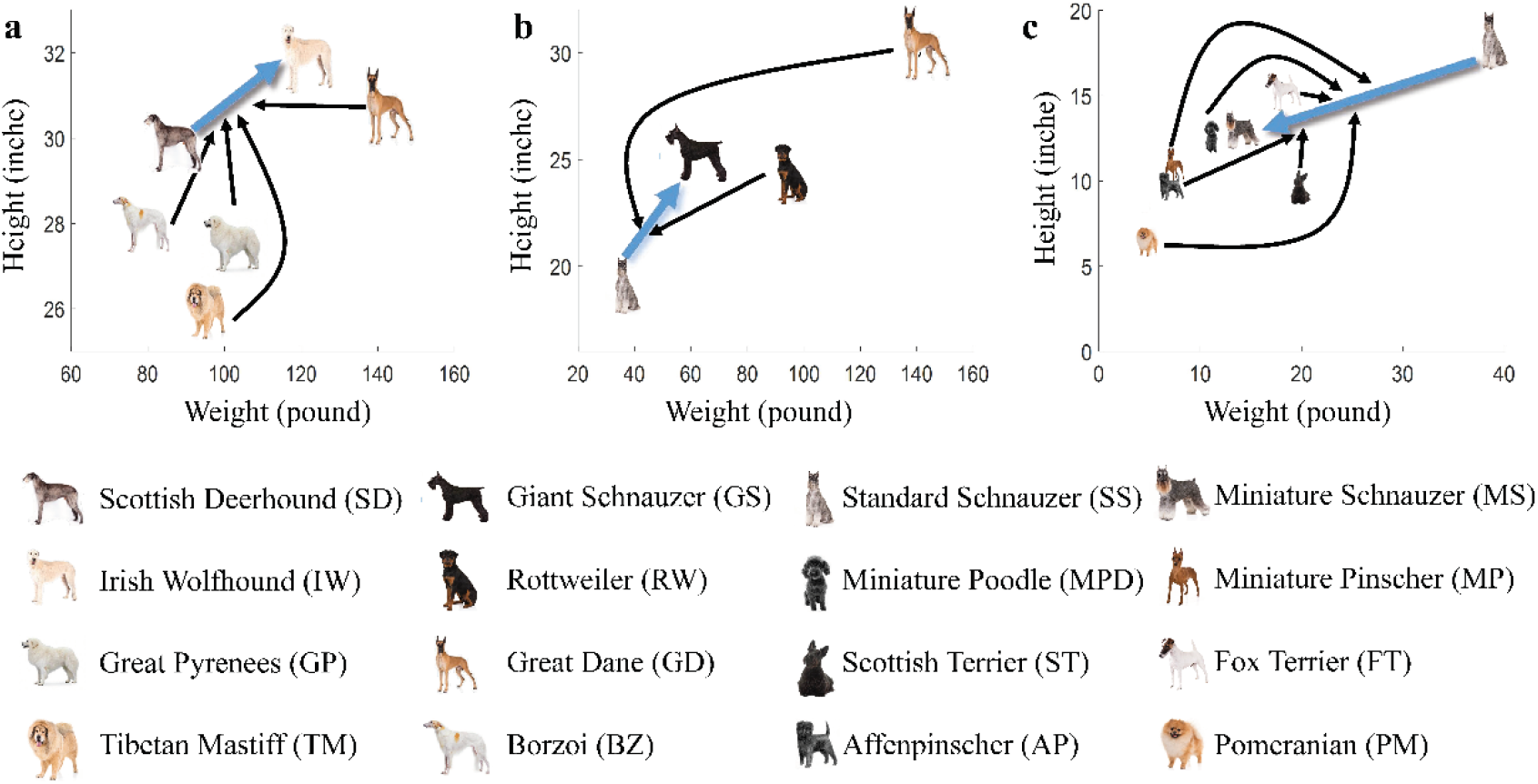
The development of three modern dog breeds aiming to change body size by interbreeding of multiple ancestral breeds. The X-axis and Y-axis are weight and height respectively. **a** In the first case several large body-size dog breeds were interbred with Scottish Deerhound to save the endangered Irish Wolfhound. **b** In the second case Giant Schnauzer was generated by using Standard Schnauzer as the baseboard breed and crossing with several breeds with larger body size. Giant Schnauzer is about 8 inches (20 cm) taller than the Standard Schnauzer. In both the first and the second cases Great Dane was used as an ancestral breed to enlarge the body size. **c** In the third case Miniature Schnauzer was generated by interbreeding several small-size dog breeds with the Standard Schnauzer being the baseboard breed as well. Miniature Schnauzer is with a height of 14 in (36 cm) or less, and Standard Schnauzer is 18-20 inches (45-50 cm).

In this study, the term “purified segment” is used to describe a specific segment in the genome of an admixed breed B that is primarily contributed by ancestral breed A (denoted as *S*_*A*→*B*_), if it is identified to be predominantly inherited from ancestral breed A with a probability greater than 0.8. These purified segments are likely the result of artificial selection aimed at retaining specific traits of an ancestral breed. Genes that overlap with *S*_*A*→*B*_ are referred to as purified genes (*G*_*A*→*B*_).

In the study we identify 288 purified segments in Irish wolfhound, 175 purified segments in Giant Schnauzer, and 185 purified segments in Miniature Schnauzer. The mean lengths of these segments are 1.47 Mb, 0.80 Mb, and 1.38 Mb respectively. Table 1 provides a summary of the purified segments in each admixed breed. The ancestry proportions of wolf are 0.52%, 2.62%, and 0.45%, respectively.

**Table 1.** Summary of purified segments for the three admixture cases.

| Descendants | Ancestral breeds | # segments | Total length (Mb) | # genes | Proportion (%) |
| --- | --- | --- | --- | --- | --- |
| Irish<br>wolfhound | Scottish Deerhound | 135 | 238.92 | 1788 | 36.99 |
|  | Great Dane | 116 | 154.56 | 1038 | 36.77 |
|  | Borzoi | 28 | 26.71 | 223 | 14.15 |
|  | Great Pyrenees | 8 | 3.38 | 39 | 9.32 |
|  | Tibetan Mastiff | 1 | 0.48 | 1 | 2.25 |
|  | Wolf | 0 | 0 | 0 | 0.52 |
| Giant<br>Schnauzer | Standard Schnauzer | 61 | 42.37 | 399 | 33.93 |
|  | Great Dane | 102 | 88.84 | 700 | 38.85 |
|  | Rottweiler | 12 | 9.06 | 45 | 24.61 |
|  | Wolf | 0 | 0 | 0 | 2.62 |
| Miniature<br>Schnauzer | Standard Schnauzer | 37 | 49.81 | 388 | 19.32 |
|  | Affenpinscher | 11 | 17.47 | 112 | 8.27 |
|  | Miniature Pinscher | 16 | 20.92 | 173 | 11.75 |
|  | Miniature Poodle | 44 | 62.56 | 401 | 21.22 |
|  | Pomeranian | 65 | 94.43 | 818 | 25.76 |
|  | Fox Terrier | 6 | 2.87 | 11 | 7.36 |
|  | Scottish Terrier | 6 | 7.15 | 129 | 5.89 |
|  | Wolf | 0 | 0 | 0 | 0.45 |

Note: purified segments are genomic regions covering with the proportion from one ancestral population $> 0.8$ . Details of the inferred local ancestral segments are shown in Supplementary Figs. 1-3.

One interesting finding is the significant presence of the Great Dane ancestry in both Irish Wolfhound and Giant Schnauzer genomes, being 36.77% and 38.85% respectively. Great Dane is the largest dog among the ancestral breeds in the two interbreeding cases. The high proportion of Great Dane in Irish Wolfhound and Giant Schnauzer highlights its critical role in the body-size enlargement of these two breeds.

### Sixty-five common genes of *G*_*GD*→*rW*_ and *G*_*GD*→*GS*_ revealing the genetic basis of large body size of Great Dane

We identify 116 purified segments of Great Dane in the genome of Irish Wolfhound, containing 1038 genes, and 102 purified segments of Great Dane in the genome of Giant Schnauzer, containing 700 genes. Interestingly, we observe a significant overlap in the gene lists of Great Dane-contributed segments between the two cases, with 65 identical genes (p-value = 2.2×10 ^-4^) scattering across 11 segments of 9 chromosomes (Table 2). The largest purified segment lies on chromosome 6, 7232952-8872321, spanning 1.64 Mb and containing 33 genes. Thus, for some of the purified regions, it is difficult to precisely identify the exact functional genes. In contrast, we can make more definitive inference for five regions including only a single gene (*LCORL, SORCS3, WDR11, FGFR2*, and *FARS2*). As shown in Fig. 3, for most of these genes, the medians of Great Dane ancestral proportions are high in the two admixture cases with a narrow range of uncertainty.

**Table 2.**
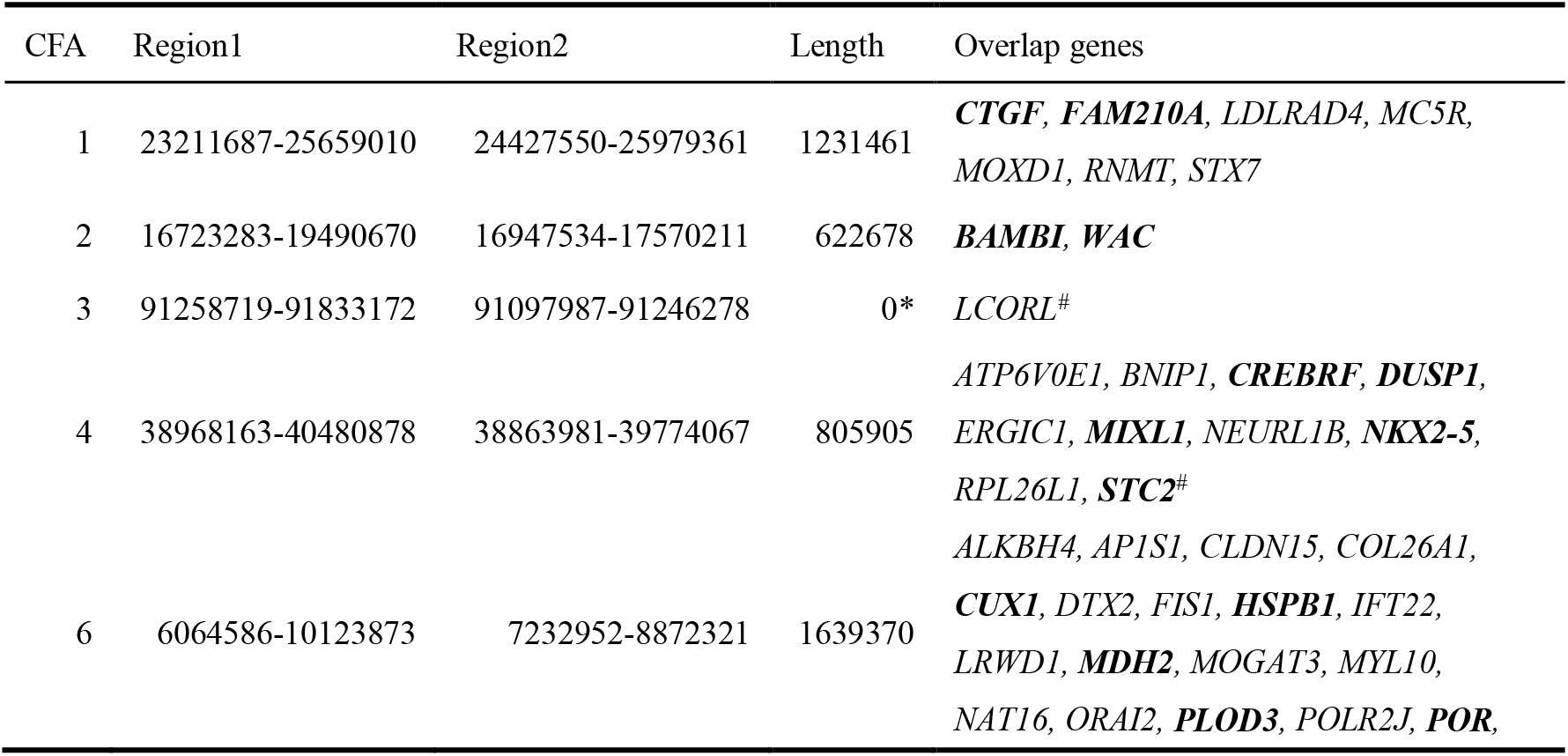
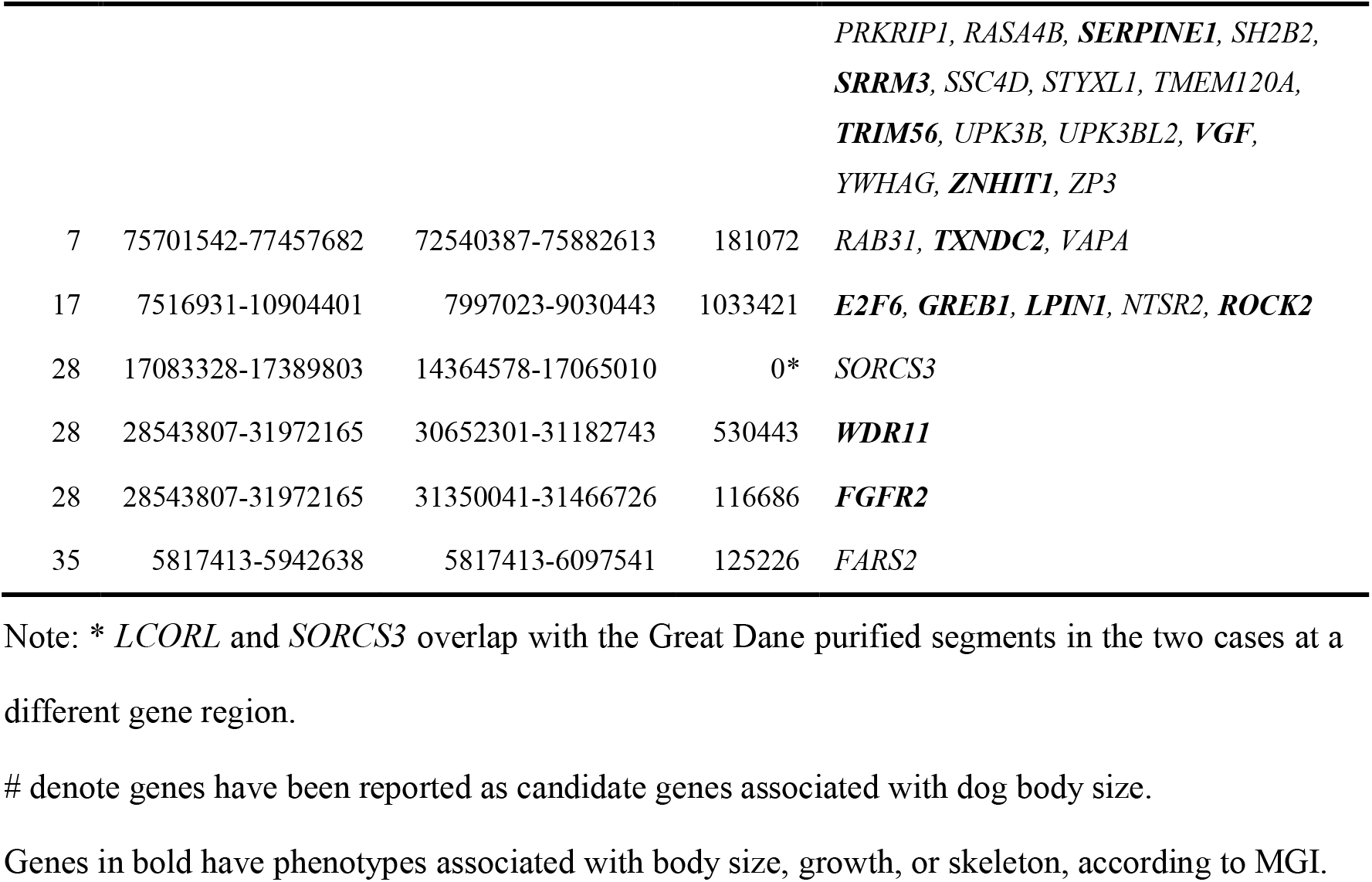
Overlapping regions of Great Dane ancestral purified segments.

**Fig. 3.**
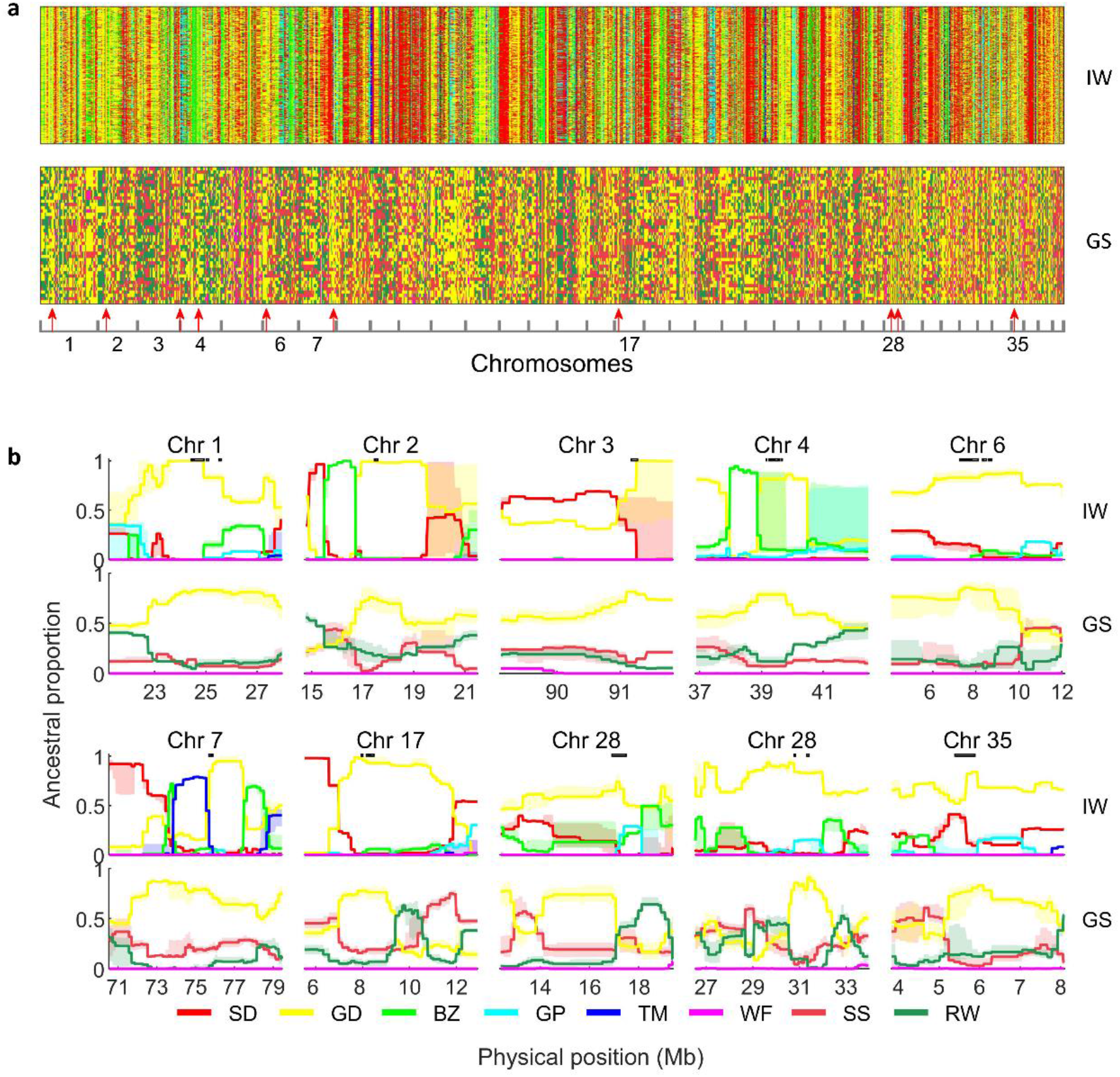
Inferring the local ancestors along the genomes of Irish Wolfhound and Giant Schnauzer identifies purified segments as candidate loci under positive artificial selection in the interbreeding process. **a** The genome-wide pattern of local ancestral segments of Irish Wolfhound and Giant Schnauzer. Each row represents a haplotype, and each column represents an SNP locus. The colors denote the ancestral from which the admixed haplotype decedent. **b** Local ancestral proportions of Great Dane purified segments in Irish Wolfhound and Giant Schnauzer. The solid lines represent the median of ancestral proportions for 100 repeats. The shadows cover the 25% - 75% quantile. The different colors represent different ancestors, as shown in the legend below. SD: Scottish Deerhound, GD: Great Dane, BZ: Borzoi, GP: Great Pyrenees, TM: Tibetan Mastiff, WF: Wolf, SS: Standard Schnauzer, RW: Rottweiler. The last two purified segments in chromosome 28 are drawn together. The location of all genes in Table 2 are shown as rectangular boxes on the top of the figures.

We calculate the empirical p-value of ancestor proportion of Great Dane for segments under artificial selection in the enlargement of Irish Wolfhound and Giant Schnauzer (see Methods for details). The histograms of local ancestor proportions in the three breeding practices are shown in Supplementary Figs. 4-6. Fig. 4 shows the Manhattan plot of p-values along the genome. A whole list of 354 genes with -log10(p) > is shown in Supplementary Table 2.

**Fig. 4.**
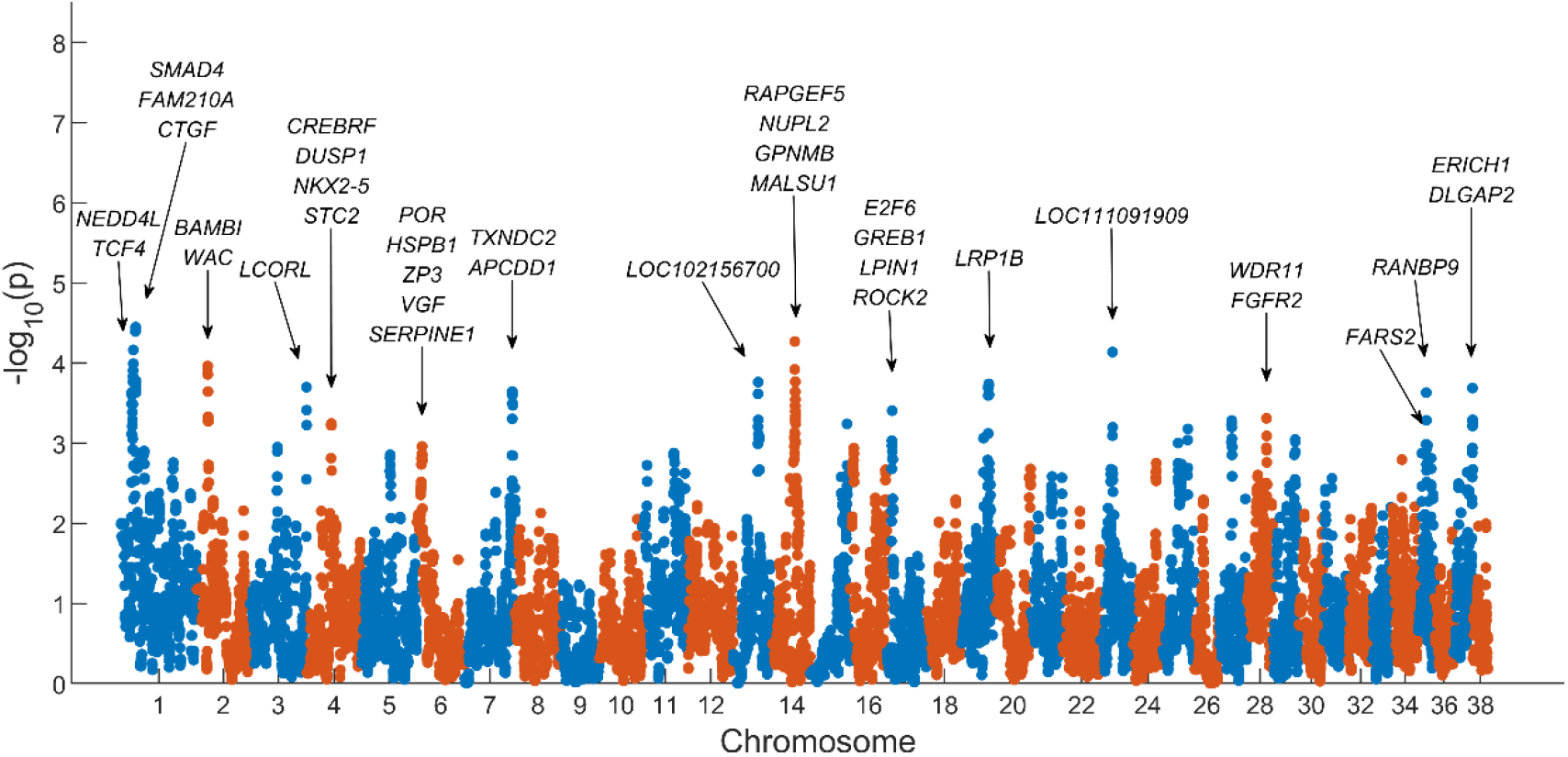
The empirical p-value of positive artificial selection of Great Dane in the body-size enlargement of Irish Wolfhound and Giant Schnauzer.

We perform Gene Ontology (GO) enrichment analysis with DAVID on the 65 overlapped purified genes contributed by Great Dane (Supplementary Table 3). Interestingly, we find that genes involved in growth-related terms are significantly overrepresented in the list. The terms include *anatomical structure formation involved in morphogenesis* (p-value = 0.0037), *regulation of developmental growth* (p-value = 0.0061), *positive regulation of developmental process* (p-value = 0.0079), and *chordate embryonic development* (p-value = 0.0091). According to the GO enrichment analysis, the five genes may play a role in controlling growth, including *FGFR2, STC2, NKX2-5, HSPB1*, and *ZP3*. In addition, 41 out of the 65 genes have a strong correlation with body height or body weight in humans GWAS (Supplementary Table 4). Twenty-six out of the 65 genes have phenotypes associated with growth, body size, limb, or skeleton, according to the Mouse Genome Informatics database (Supplementary Table 5). Two genes are reported as candidate genes responsible for dog body size in former studies (*LCORL* ^8, 10, 13^ and *STC2* ^8, 14^).

### Candidate artificial selection regions with single-gene

We further explore the function of the five single genes contained in the purified segments contributed by Great Dane. For purified segments that contain multiple genes, a detailed illustration can be found in Supplemental materials.

*FGFR2* locates on a Great Dane purified segment of Chromosome 28 (Fig. 3B). Essential roles of Fibroblast growth factor (FGF) and their binding receptors (FGFR) signaling in skeletal development and diseases have been reported in humans and mouse models ^22^. *FGFR2* is expressed in condensing mesenchyme of early limb bud, and later appears as the marker of prechondrogenic condensations. In developing bone, *FGFR2* is predominantly localized to perichondrial and periosteal tissues, and weakly to endosteal tissue and trabecular bone. *FGFR2* is intensely expressed in the cartilage of the cranial base and growth plate. In cranial sutures, *FGFR2* is mainly expressed in osteoprogenitor cells and differentiating osteoblasts. The expression pattern of *FGFR2* indicates its important role in skeleton development.

Fgfr2 mutant mice exhibit various phenotypes. Fgfr2^+/S252W^ mutant mice have smaller body size. Fgfr2^+/P253R^ mice have growth retardation of the synchondroses of cranial base and growth plates of the long bones with decreased proliferation of chondrocytes, which may be responsible for the smaller body size and shortened cranial base in Fgfr2^+/P253R^ mice. Conditional deletion of Fgfr2 in mesenchymal condensations of mice via Dermol-Cre results in skeletal dwarfism and decreased bone density.

Fgfr2IIIc^−^/^−^ mice also exhibit dwarfisms, reduced growth of the skull base and axial, as well as appendicular skeletons, which is associated with decreased proliferating chondrocytes and hypertrophic zone in these endochondral bones. This leads to premature loss of skull base sutures and smaller-than-normal long bones and vertebrae (see details in the review article ^22^).

*WDR11* is a single gene locates on another purified segment of Chromosome 28. It is in the Indian hedgehog (Ihh) signaling pathway. Once the limb bud is patterned and the skeletal tissues have differentiated, the regulation of the long bone growth plates greatly contributes to the overall size of the adult limb. Indian hedgehog (Ihh) signaling positively regulates cell proliferation within the growth plates of the long bones ^23^. When Ihh signaling is inactivated, through null Ihh or mutations in Ihh transducers or effectors, the resulting mammalian limbs are severely shortened ^24, 25, 26^. In contrast, overexpressing Ihh in the developing chick limb through viral transfection resulted in increased limb length ^27^. These effects on limb size are generally tied to altered Ihh signaling during the processes of chondrocyte proliferation and differentiation and osteoblast differentiation in the growth plates in the long bones ^28^.

The Wdr11 deficient mice exhibited multiple phenotypes such as increased body mass index, obese, postnatal growth retardation, abnormal skeleton morphology, decreased bone mineralization, and short limbs ^26^. Mutants of *WDR11* also have been demonstrated to be causative for human growth retardation ^29^.

*LCORL* locates on Chromosome 3 and is a transcription factor famous for controlling body size. Several studies have associated *LCORL* locus with domestic dog body size ^8, 10, 13^. The association also exists in multiple domestic mammals. Expression levels of *LCORL* are associated with body size within and across horse breeds ^30^. *LCORL* is under natural selection in several large-sized Pakistani goat breeds ^31^. Whole-genome analyses also identified the association of *LCORL* with body size in cattle ^32^. Selective sweep analyses revealed strong signatures of selection at *LCORL* in the domestic pig genome, harboring quantitative trait loci related to the elongation of the back and an increased number of vertebrae ^33^. *LCORL* is also identified being associated with human height in GWAS ^34, 35, 36^.

*FARS2* is from a purified segment of Chromosome 28, which encodes the mitochondrial phenylalanyl tRNA synthetase. Mutations in *FARS2* cause defects in mitochondrial protein synthesis and result in various mitochondrial disease phenotypes.

*FARS2* deficiency in Drosophila leads to developmental delay by aberrant mitochondrial tRNA metabolism ^37^. *FARS2* variants in humans also result in global developmental delay ^38, 39^, and is significantly associated with human weight ^40^.

### Genetic architecture of body-size enlargement of Irish Wolfhound

In addition to the common purified genes identified in Irish Wolfhound and Giant Schnauzer, we also investigate the biological functions of the 1038 genes in **G**_**G**D→IW_ by performing Gene Ontology (GO) enrichment analysis with the package DAVID (Supplementary Tables 6 and 7). The two terms, growth (7 purified genes with a p-value of 0.059) and developmental growth (6 purified genes with a p-value of 0.062) are among the top terms in Supplementary Table 7.

Among the seven genes in the growth term, *TTC8, GJA1, CTNNB1*, and *FGF1* are unique in the Irish Wolfhound breed. *TTC8* is associated with Bardet-Biedl syndrome in humans, characterized by a short neck, short stature, and obesity ^41^. *TTC8* mutant mice exhibit obese and decreased body size ^42^. *GJA1* links with various phenotypes related to body size, growth, limbs, and skeleton in humans and mice ^43, 44, 45, 46, 47, 48, 49^. *CTNNB1* is involved in the classical Wnt/β-catenin signaling pathways, and associated with the development of osteoblasts ^50^. *CTNNB1* is also associated with various phenotypes related to body size, growth, limbs, digits, and skeleton in humans and mice ^50, 51, 52, 53, 54, 55^. *FGF1* is a member of the FGF/FGFR signaling pathway, which plays an essential role in skeletal development and homeostasis, including the *FGFR2* gene we discussed above. However, mice lacking *FGF1* show normal skeleton phenotypes 22.

In the Irish Wolfhound genome, several purified genes contributed by Scottish Deerhound have been identified to be associated with dog body size. These genes include *HMGA2, GNS, RASSF3*, and *SOX9* ^9, 15, 56^. In mice, mutations in *HMGA2* result in the pygmy phenotype ^57^, characterized by aberrations in adiposity and disrupted growth leading to dwarfism. Sox9 is a member of the SRY-related HMG-box family of transcription factors, which effect limb size through their regulation of chondrogenesis ^58^. During embryonic limb development, Sox9 is considered the master chondrogenic factor required for differentiating mesenchymal precursor cells into chondrocytes ^58, 59^. Sox9 then works in concert with Sox5 and Sox6 to drive the differentiation and proliferation of chondrocytes ^58, 59^. Activating mutations in Sox9 in mice results in a long limb phenotype ^60^, while inhibiting mutations in the same gene results in short limb phenotypes ^61^. These observations highlight the importance of this family of transcription factors on the regulation of growth during limb development. *GNS* avian ortholog (QSulf1) regulates WNT signaling during embryogenesis in myogenic somite progenitors ^62^.

### Genetic architecture of body-size enlargement of Giant Schnauzer

We investigate the biological functions of the 700 *G*_*GD*→*GS*_ by performing Gene Ontology (GO) enrichment analysis with the package DAVID (Supplementary Tables 8 and 9). Several top terms related to growth, including regulation of growth (10 purified genes with p-value 7.1×10 ^-5^), regulation of developmental growth (7 purified genes with p-value 6.2×10 ^-4^), and growth (9 purified genes with p-value 0.0010). These terms contain 11 genes including *GHSR, TGFB1, PROC, TFRC, BCL6, IFNG, HSPB1, PRL, ZP3, NKX2-5*, and *FGFR2*. Among the 11 genes, *HSPB1, ZP3*, and *NKX2-5* are also purified genes in Irish Wolfhound. *GHSR* is primarily expressed in the brain and pituitary gland, where it mediates ghrelin’s effects on food intake and growth hormone secretion. Mutations in this gene are associated with autosomal idiopathic short stature ^63, 64, 65^. *TGFB1* links with decreased body weight and height in mice ^66^. *TGFB1* is one of the key regulators of feed efficiency in beef cattle ^67^. *TFRC* is associated with embryonic growth retardation in mice ^68^. Mutations in *BCL6* are associated with reduced body size and postnatal growth retardation in mice ^69, 70, 71^. *IFNG* is associated with phenotypes of decreased body size and postnatal growth retardation in mice ^72^, and short stature in humans ^73^.

Except for genes in the above growth-related terms, we find several other *G*_*G*D→*GS*_ genes that may be responsible for dog body size, including *NPR2* and *FGF8. NPR2* has been reported as a candidate gene for size-related traits of dog ^56^. Loss-of-function mutations in *NPR2* cause dwarfism in humans ^74^. In contrast, gain-of-function mutations in *NPR2* cause Overgrowth syndrome ^75, 76^. *NPR2* is also associated with the phenotypes of growth, body size, limbs, and skeleton in mice ^77^. *FGF8* is expressed throughout the apical ectodermal ridge (AER), indicating its essential role in limb development ^78, 79, 80^. Conditional deletion of Fgf8 in the developing forelimb AER results in severe forelimb deformity, including the absence of radius and first digit ^81, 82^.

### Genetic architecture of body miniature of Miniature Schnauzer

In the development of Miniature Schnauzer, six breeds with small body size were interbred to contribute functional loci for miniature (Fig. 2c, Table 1). Genes on purified segments from these ancestral populations in the Miniature Schnauzer genome are thus candidate loci of miniature.

1. Genes contributed from Affenpinscher: *FGF4* is involved in the biological process of limb development, limb morphogenesis, and skeletal system morphogenesis. Vertebrate limb development largely depends on signals from apical ectodermal ridge (AER). *FGF4* is first expressed in the developing murine forelimb bud during limb development at E10.0. Its expression is strongest in the posterior AER at E10.5–11.0. *FGF4* provides mitogenic and morphogenic signals to regulate normal limb development ^83, 84^. An expressed fgf4 retrogene is associated with breed-defining chondrodysplasia and height in domestic dogs ^8, 10, 85^. *LEP* and *BRCA2* are involved in the biological process of developmental growth. *LEP* is a skeletal growth factor and is essential in the regulation of energy balance and body weight control ^86^. *LEP* mutated mice exhibit significantly elevated body weights and decreased length of long bone ^87^. *BRCA2* mutant mice weigh half as much as wild-type mice ^88^. *FGF3* knock-out mice show a short, dorsally curled tail, caudal vertebrae, and smaller body ^89^.
2. Genes contributed from Pomeranian: *TFAP2B* is involved in the biological process of limb development and limb morphogenesis. *TFAP2B* mutant mice exhibit postnatal growth retardation ^90^. *BCL6* is involved in biological process growth. Interestingly, *BCL6* is also implicated in the enlargement of the Standard Schnauzer.
3. Candidate genes contributed from Miniature Pinscher: Fibroblast growth factor receptor 3 (*FGFR3*) is expressed in the growth plate of endochondral bones and serves as a negative regulator of linear bone elongation. Activating mutations severely limit bone growth, resulting in dwarfism, while inactivating mutations enhance bone elongation and overall skeletal size. Gain-of-function mutations in both humans and mice *FGFR3* result in achondroplasia characterized by a short limb phenotype ^91, 92, 93, 94^, while knock-out of *FGFR3* in mice produces a long limb phenotype ^95, 96^. Furthermore, the knock-out of FGFR3 has been directly tied to increased Ihh and BMP signaling within the elongating skeletal tissue in mice ^96^. Although a previous study shows that the *FGFR3* gene is conserved across nine dog breeds representing a spectrum of skeletal size ^97^, we find genetic diversity within the *FGFR3* region using a larger dataset of ∼500 breeds (including 223 village dogs) and 1629 whole genome sequencing samples (Supplementary Fig. 7). In addition, *FGFR3* is proposed to be the gene responsible for the foreshortened limbs in Dachshunds in a study using selective sweep mapping ^98^.
4. Candidate genes contributed from Fox Terrier: *FGF5* is involved in the biological process of response to growth factor. *FGF5*-lacking mice show normal skeleton phenotypes ^22^.

### Breed-specific purified genes in the bone development-related pathways

In the above analysis of three breeding cases we identify numerous breed-specific purified genes that potentially play a role in the enlargement or miniaturization of body size under artificial selection. These genes are significantly enriched in signaling pathways related to bone development, including FGFR, BMP signaling, TGF-beta, and Wnt (Fig. 5, Supplementary Tables 10-12).

**Fig. 5.**
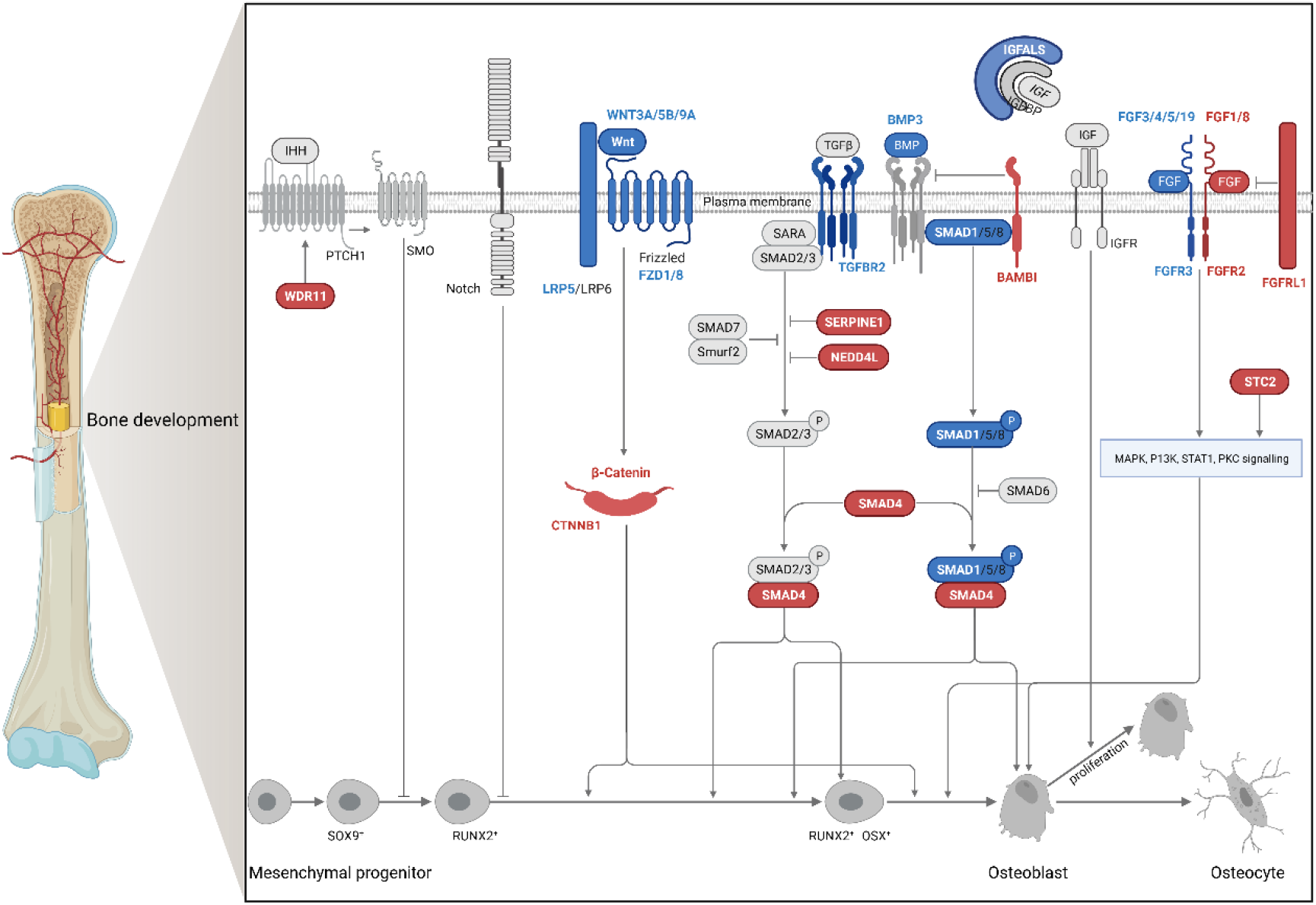
Genes under breed-specific artificial selection are enriched in the bone development-related signaling pathways. Genes marked in red contribute to the enlargement of the baseboard breed, and genes marked in blue contribute to the miniaturization of the baseboard breed.

Interestingly, when we examine a subset of 3,849 genes associated with body height in a human GWAS with a p-value <10-15 ^99^, we find a significant overlap with our gene sets (p-value=0.0021 for 354 genes with -log10(p-value) <2.5, and p-value=0.0030 for 65 genes purified from the two Great Dane ancestors). Furthermore, the human height-associated genes also exhibit enrichment in the bone development signaling pathways, including FGFR, BMP, TGF-beta, and Wnt (Supplementary Table 13). These findings suggest the critical roles of these pathways in determining body size, and the common mechanism underlying body size in both dogs and humans.

We further check the effect sizes in human GWAS of those variants with the most significant p-values in the GAMA analysis. The mean absolute effect size for 6135 human GWAS genes is 0.0115, while the value is 0.0157 for the 22 artificial selection genes in dogs (22 out of 65 genes overlap with human GWAS significant gene sets) and 0.0132 for 112 artificial selection genes in dogs (112 out of 354 genes overlap with human GWAS significant gene sets). The effect size of the 22 dog artificial-selection genes is 36.52% higher than that of the European ancestry significant genes (p-value 0.0425), and the mean effect size for the 112 gene set is 14.78% higher than the European genes (p-value 0.0836). These findings suggest that artificially selected genes related to body size change in dogs tend to exhibit higher effects in human GWAS, aligning with the expectation that artificial selection typically acts on major genes with substantial effects. However, it is worth noting that the differences in effect sizes are not pronounced, potentially due to the presence of different causal mutations in the corresponding genes of humans and dogs.

### Population differentiation between large dog breeds and small dog breeds

To compare with the GAMA analysis, we also carry out a genome-wide selective sweep scan using population differentiation. Fst (fixation index) as a measure of population differentiation is calculated for each SNP between the two groups: the fourteen small and the nine large dog breeds (see Methods for details). The top 10 significant signals and the candidate genes are shown in the genome-wide Manhattan plot in Fig. 6. Among these genes, the most significant one, *IGF1*, which encodes insulin-like growth factor 1, has been widely reported as a major contributor to body size miniature in all small dogs. *SMAD2* is a transcription factor known to transduce signals from members of the transforming growth factor beta (TGF-beta) superfamily ^100, 101^. A deletion identified proximal to *SMAD2* demonstrates the cis-effect to alter this gene’s expression in developmental processes such as myogenesis, chondrogenesis, or osteogenesis ^102, 103, 104^. *IRS4* is reported to explain the body size variance in “large breeds” with an SBW >41 kg ^10^ but not the variance between large and small breeds. *CHRDL1* encodes an antagonist of bone morphogenetic protein 4. *NR1D2* may play a role in circadian rhythms, and carbohydrate and lipid metabolism. *SYNE1* encodes a spectrin repeat containing protein expressed in skeletal and smooth muscles, and peripheral blood lymphocytes. *LCORL* is associated with skeletal frame size and adult height. *RSPO2* is involved in the GO term Limb morphogenesis. The other two genes, *ATP23* and *BANF2*, have less evidence to mediate the body size of dog breeds.

**Fig. 6.**
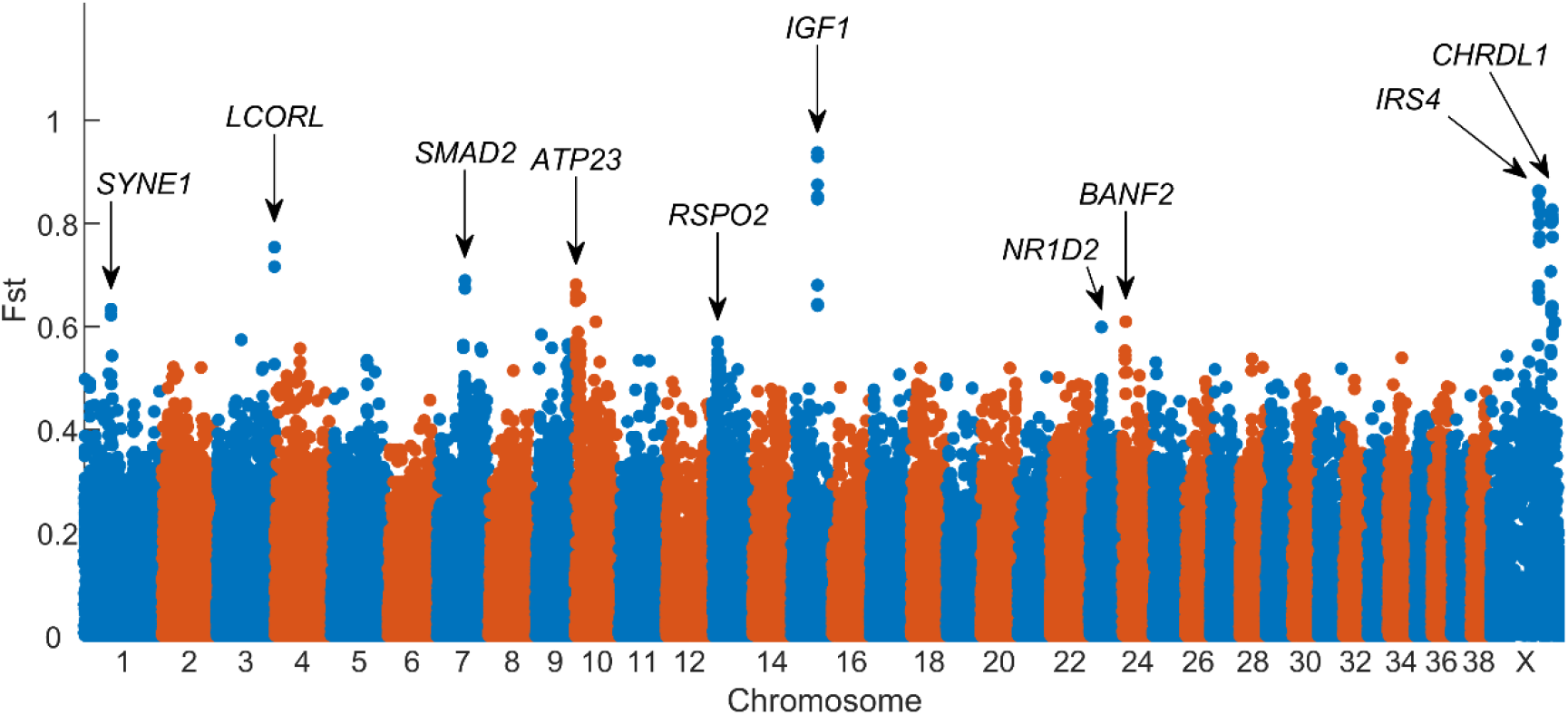
The top 10 signals found by Fst analyze between 14 small and nine large dog breeds.

Remarkably, there is very limited overlap between the two sets of body-size related genes identified by Fst and GAMA methods (e.g., *LCORL*). Fst identifies common genes shared by multiple small or large dog breeds, while GAMA detects breed-specific genes underlying the phenotypes by constructing genetic ancestry along the genome. By combining the two approaches, a more comprehensive understanding of the genetic architecture underlying body size traits can be achieved.

## Discussion

The vast phenotypic diversity of domestic dog breeds has long been recognized as a unique portal into the genetic architecture of phenotype. Here, we present a new method, GAMA, for accurately inferring genomic segments with different ancestral origins from multiple admixture. Simulations demonstrate that GAMA outperforms existing methods in inferring local genomic ancestry, especially for long-term and multi-way admixtures. Accurate inference of the mosaic of ancestral segments for domestic animals and plants with complex interbreeding history allows decoding genetics underlying the complex traits with ancestor-specific origins.

We apply GAMA to three dog interbreeding cases with body size enlargement and miniaturization. We find multiple purified genes as candidate genetic loci under artificial selection in the breeding process. Interestingly, these genes are significantly enriched in the meaningful biological process, such as growth and development. In interpreting the signatures of artificial selection identified, we leverage information about gene function from other species. Many candidate genes show evidence implicated in body size, growth, and skeleton phenotypes in mice and other species. Previously identified genes associated with dog body sizes, such as *LCORL, STC2, NPR2*, and *FGF4*, are validated in the study. Multiple previously unidentified genes, such as *FGFR2, WDR11*, and *FARS2*, are also found to be presented in highly purified segments in admixed dog breeds. The results indicate the applicability and efficiency of GAMA for investigating complex traits with ancestry-specific origins in the domestic animals with complex interbreeding history.

Like traditional association studies and selective sweep analysis, a paucity of recombinants in dog breeds can provide only a coarse scale mapping of the potential causative loci using our method. Analyzing several admixed dog breeds sharing the same ancestors can help narrow down the confidence intervals of candidate gene locations. The hundreds of admixed dog breeds provide an excellent model for delineating the phenotypic effects in dogs utilizing our method.

Despite the novel insights provided by the new approach, several limitations remain for further study. The performance of GAMA in inferring local ancestries is affected by the determination of ancestors implicated in the breeding process and the sufficiently large samples representing the genetic diversity of reference populations.

However, interbreeding records of many dog breeds may be obscure, considering the long breeding history of modern dog breeds initiated in the Victorian era. Although the accumulation of dog genome data is fast, some ancestral populations still have a small genome sample size, leading to a bias toward underestimating the ancestral composition. Additionally, if the causal mutation is shared simultaneously by several ancestors, we may miss the causal mutations by identifying purified segments. An example is *IGF1* in the miniaturization of Standard Schnauzer. The composing proportions of *IGF1* from Affenpinscher, Miniature Pinscher, Miniature Poodle, Pomeranian, Fox Terrier, Scottish Terrier, and Wolf are 0, 0.0069, 0.0141, 0.4911, 0.0001, 0.4256, and 0, respectively. No ancestor-specific purified segments are identified around *IGF1*. For such cases, a population differentiation analysis for a merged group of multiple breeds can be helpful.

## Acknowledgments

The work was supported by the Science and Technology Innovation 2030-Major Project (2022ZD04017), the China National Natural Science Foundation (Grant No. 32370669) and the Strategic Priority Research Program of the Chinese Academy of Sciences (XDB38040200). Genomic variation data was produced by the Dog10K Project, an international collaboration to advance canine genetics. Genome sequencing was supported by National Science and Technology Innovation 2030 Major Project of China (2021ZD0203900) and the National Key R&D Program of China (2019YFA0707101).

## Method

### GAMA model for inferring genetic ancestry segments in admixed individuals

The genome of an admixed individual resembles a mosaic of ancestry blocks that are inherited from different ancestral populations, with changes in ancestry occurring at recombination points. The mosaic pattern depends on time (in generations) since admixture, recombination rate, and ancestry proportions. We proposed a novel method, Genomic Analysis of Multiple Admixture (GAMA), to infer local ancestry segments from multi-way admixture along the genome. The model contains three free parameters, the number of generations since admixture *λ*, ancestry proportions *π* of each contributor, and error rates for the haplotypes *α*.

### Input data

GAMA requires phased haplotypes from unrelated individuals for ancestral populations and admixed individuals, and genetic distances between adjacent loci. The genetic distances can be obtained from the genetic map or approximated using global recombination rate. The phased haplotypes are subsequently divided into *L* successive local fragments that contain the same number of *k* SNPs, with *H* = {*h*_1_, *h*_2_, …*h*_*L*_ }. The local haplotype distance is denoted as *D* = {*d*_1_, *d*_2_, …*d* _*L*−1_}, where *d*_*l*_ represents the genetic distance between the center of locus and *l* +1.

### Hidden variables

The sequence of the hidden variables is denoted as *Z* = {*z*_1_, *z*_2_, …*z*_*L*_ }, *z*_*l*_ ∈{1,…, *N*}, where *z*_*l*_ = *n* represents that chromosome fragment containing local haplotype is inherited from ancestral population *n*, and *N* represents the number of the ancestral populations.

### Transition matrix

The transition probability of hidden states is determined by genetic distance, admixture generations, and admixture proportions. The transition matrix at locus *l* is

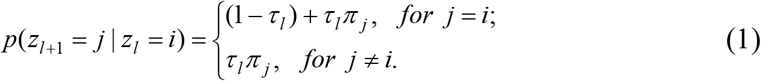

Where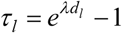is the probability of recombination occurring between locus *l* and *l* +1, *λ* is admixture time, _*d*_*l* is genetic distance between locus and *l* +1, and *π j* is the ancestry proportion of population *j*. Considering the low recombination rate, we simplify the transition probability by ignoring the repetitive recombination between the adjacent local haplotypes.

### Emission probability

The emission probability is chosen to be the haplotype frequencies of a given ancestral population and modified by haplotypes error rate of the admixed population. The haplotypes error rate take into account the effects of mutation, sequencing error, or phasing error. Suppose that the local haplotype segment contains *k* SNPs, the possible number of observed haplotypes is 2^*k*^. The optimized setting of the bin size *k* is discussed in the Supplementary Materials. Any haplotypes are possible to be observed from any ancestral states under the non-zero haplotypes error rate. Suppose the haplotype frequencies of ancestor *n* at locus *l* are the emission probability is *f*_*i*_, *i* = 0,1,…,2^*k*^ − 1,

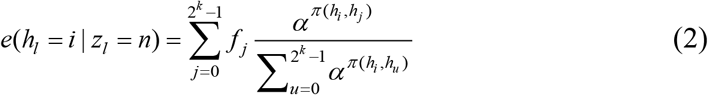

where *h* denotes the local haplotype, *π* (*h*_*i*_, *h*_*j*_) is the total nucleotide difference between haplotypes *h*_*i*_ and *h* _*j*_, and *α* is the error rate as we mentioned above. Error rate represents the probability of SNP with allele type different from the inherited ancestor. If we set *k*=1 and *α*=0, the emission probability will be

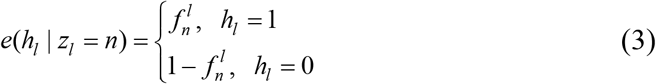

Where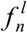is the alternative allele frequency of ancestor population *n* at locus *l*.

### Forward-backward algorithm

Let *θ* = (*π, λ,α*). Denote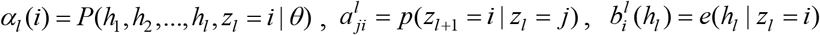 and the forward functions,

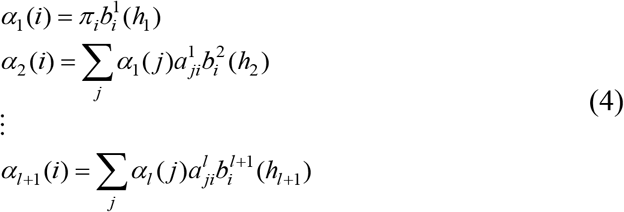

Denote*β*_*l*_ (*i*) = *P*(*h*_*l* +1_, *h*_*l* +2_, …, *h*_*L*_, *z*_*l*_ = *i* | *θ*), and the backward functions,

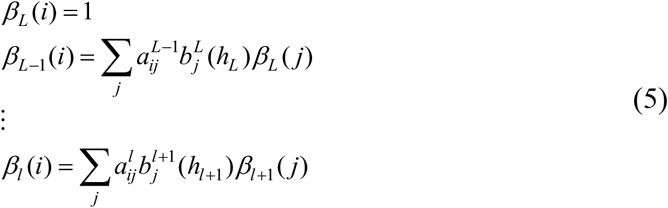

The probability of observations given parameter can be written as,

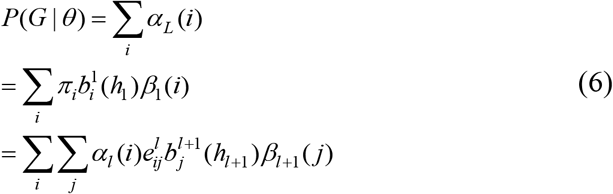

Where 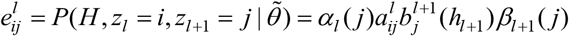

### Expectation Maximization algorithm for inference of parameters

An expectation-maximization (EM) algorithm was applied to infer free parameters.

E-step:

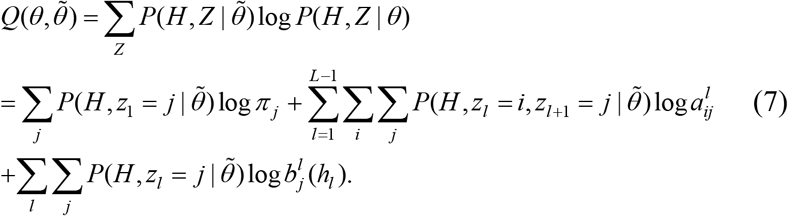

Where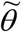 are the current estimation of the parameters.

M-step:

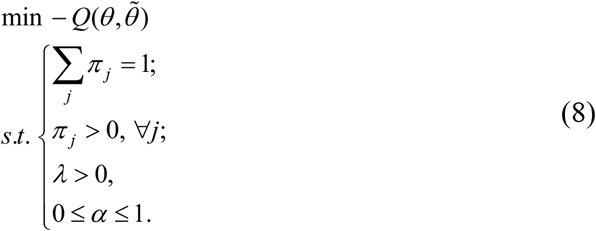

For convenience we denote

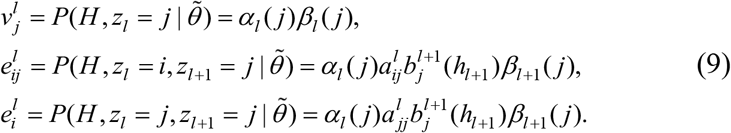

Then the above M-step can be written as minimize

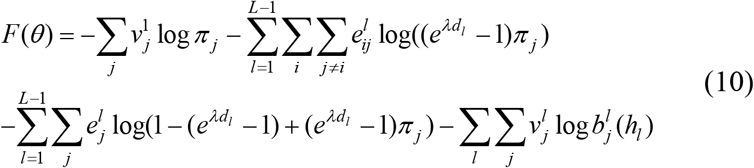

### Viterbi algorithm for inference of hidden states

The Viterbi algorithm was applied to infer hidden states.

For ⩝*i* ∈1,2,…,*N, l* ∈1,2,…,*L*, define,

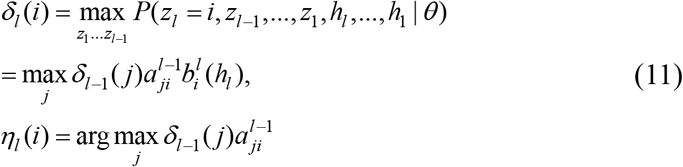

The inputs of Viterbi algorithm are *H* = (*h*_1_,…, *h*_*L*_), *θ* = (*π, λ,α*), and the output is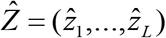

Step1: For *i* ∈1,2,…,*N*,

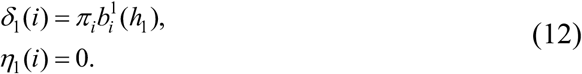

Step2: For *l* = 2,…,*L*,

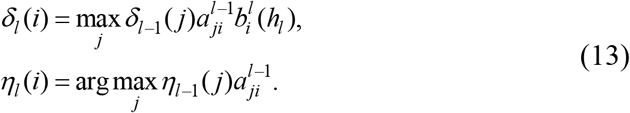

Step3:

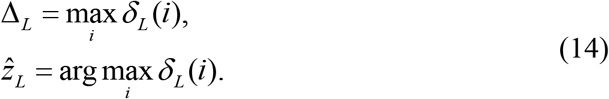

Step4: For *l* = *L* −1, *L* − 2,…,

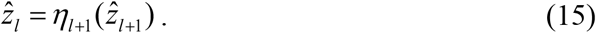

And thus, we obtain the estimated hidden states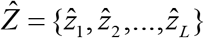

### Posterior probability of ancestral states

We estimate the marginal posterior probability that an allele inherited from a specific ancestral population,

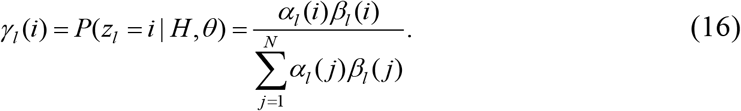

The local ancestral proportion of ancestor *i* at locus *l* is the mean of *γ* _*l*_ (*i*) across all admixture population samples.

### Significance testing of non-positive artificial selection

We state the null hypothesis that ancestor *A* at locus *l* is not under positive artificial selection in the breeding process.

Denote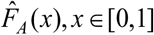 as the empirical cumulative distribution function of local ancestral proportions of ancestor *A*. If under positive artificial selection, the local ancestral proportion of ancestor *A* at locus *l* tends to increase. The empirical probability of rejecting the null hypothesis is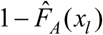, where *x* is the local ancestral proportion of ancestor *A* at locus *l*. Suppose *A* is involved in multiple independent breeding practices. We state the null hypothesis that ancestor *A* at locus *l* is not under positive artificial selection in all the breeding practices. Then the probability of rejecting the null hypothesis is the product of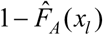 in all cases.

### The alternative way for inferring admixture proportions and admixture time

As we tested, inference of all parameters from the EM step is time-consuming and inaccurate in the estimation of admixture time. Alternatively, we only estimate the error rate *θ* = (*π, λ,α*) from the EM step. After every iteration of the EM and Viterbi algorithm steps, we infer the admixture proportion *π* and admixture time *λ* from the current estimation of local ancestry segments. Admixture proportion *π* is calculated from the current inference of hidden variables directly, and the admixture time *λ* is estimated by the co-ancestry curve method. Co-ancestry curve plots the frequencies that a pair of haplotype chunks separated under a given distance for each respective population ^105^. Here we applied the co-ancestry curve of a single admixture event,

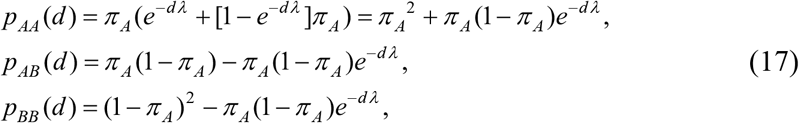

where *d* denotes genetic distance, *λ* denotes the time in generations since admixture, and *π* _*A*_ denotes the ancestral proportion of population *A*.

### Data collection and filtering

We merge whole genome sequencing samples from Dog10K project and CHIP DNA samples. For the study of Irish Wolfhound, the collected sample include 267 Irish Wolfhound, 14 Scottish Deerhound, 29 Great Dane, 22 Borzoi, 20 Great Pyrenees and 6 Tibetan Mastiff. For the study of Giant Schnauzer, the collected sample include 21 Giant Schnauzer, 14 Standard Schnauzer, 29 Great Dane, and 101 Rottweiler. For the case study of Miniature Schnauzer, the collected samples include 73 Miniature Schnauzer, 15 Standard Schnauzer, 4 Affen Pinscher, 31 Miniature Pinscher, 19 Miniature Poodle, 16 Pomeranian, 8 Fox Terrier, and 32 Scottish Terrier. We filtered outlier samples according to principal component analysis (Supplementary Figs. 8-10). We removed 2 samples from Standard Schnauzer, 1 Scottish Deerhound, 4 Miniature Pinscher, and 1 Scottish Terrier. The remaining number of samples from these four populations are 13, 13, 27, and 31 respectively. We also included Wolf as ancestral population in the three admixture cases. The number of wolf samples is 95. We applied Beagle 5.1 to phase the genotype data.

### Running GAMA

We set bin size of 10 and applied genetic map. All haplotypes from the same admixed population and the same chromosome are run together to estimate the common parameters of admixture timing, admixture proportion, and error rate. We applied all available genomes of each reference population to infer the local ancestor of admixed populations. To reduce artificial bias that may be caused by the different number of reference genomes of each ancestral population, we randomly select same number of reference genomes for each ancestral population and run the program for 100 times. Regions with the median line of local ancestral proportion of 100 repeats close to the ancestral proportion of the whole data and with a narrow confidence interval are regarded as robust.

### Running Mosaic

Mosaic was downloaded from https://maths.ucd.ie/~mst/MOSAIC/, and run with the default setting of parameters ^21^.

### Gene ontology enrichment analysis

Gene ontology enrichment analysis for purified gene sets using the Database for annotation, visualization and integrated discovery (DAVID) ^106^, https://david.ncifcrf.gov/.

### Population structure analysis

SmartPCA program from the EIGENSOFT package (version 6.1.4) was used to perform principle component analysis on the dog breeds of the three breeding cases ^107^.

### Phasing

Beagle version 5.1 was used to infer the haplotype phase with default parameters ^101^.

### Fst calculation

We used Weir and Cockerham formula ^108^ to calculate Fst between large dog breeds and small dog breeds. 26 Chihuahua, 4 Toy Fox Terrier, 16 Pomeranian, 16 Yorkshire Terrier, 4 Japanese Chin, 13 Chinese Crested, 16 Italian Greyhound, 23 Pekingese, 37 Shih Tzu, 67 Cavalier King Charles Spaniel, 38 Border Collie, 70 Miniature Schnauzer, 37 Jack Russell Terrier, 31 Boston Terrier, total samples: 398. 16 Giant Schnauzer, 22 Akita, 34 Bernese Mountain Dog, 20 Great Pyrenees, 11 Bullmastiff, 267 Irish Wolfhound, 37 Saint Bernard, 29 Great Dane, 22 Mastiff, total samples: 458.

### Simulation of genomic data under different interbreeding history

We adopted Canis lupus familiaris chromosome 1 (CanFam3.1) in simulations with 6,611 SNPs scattering in the 122.68 Mb region of the whole chromosome (the same as the actual data we analyzed). We used a uniform recombination rate of 1×10^−8^ per bp per generation. The admixture time was set to 5, 10, 20, 50, and 100 generations, and error rates of 0, 0.02, and 0.05. Sixty simulation with different parameter setting combination were carried out and for each 100 samples were generated.

Simulation was implemented along chromosome. The first SNP was randomly selected from one ancestral population according to ancestry proportions and one haplotype was randomly selected as ancestral haplotype given the ancestral population. The allele type of the ancestral populations was the same as the selected haplotype. For the n-th SNP, we calculated the recombination probability between n-th SNP and (n-1)-th SNP according to admixture timing and recombination rate. If there is no recombination event, we use the same selected haplotype as in n-th SNP; else, we will randomly select another haplotype as the ancestral haplotype. We repeated the above process for each simulation 20 times to obtain an admixture population with 20 haplotypes.

