## Supplemental tables for "Decoding genetic architecture of dog complex traits by constructing fine-scale genomic ancestry of admixture": SupplementaryInformation.docx

**1. The ancestry of genomic segments of admixed breeds inferred by GAMA**

We inferred the local ancestry of genomic segments of Irish Wolfhound, Giant Schnauzer, and Giant Schnauzer using GAMA. Supplementary Figs. S1-S3 show the local ancestry along the genomes of Irish Wolfhound, Giant Schnauzer, and Giant Schnauzer, respectively. In the line charts, the x-axis represents the physical positions along the chromosome, and the y-axis is the local ancestral proportions. In the mosaic figures, the x-axis represents the physical positions on the chromosome, and the y-axis corresponds different haplotypes of the sample. The colors denote different ancestors of genomic segments of the admixed individuals.

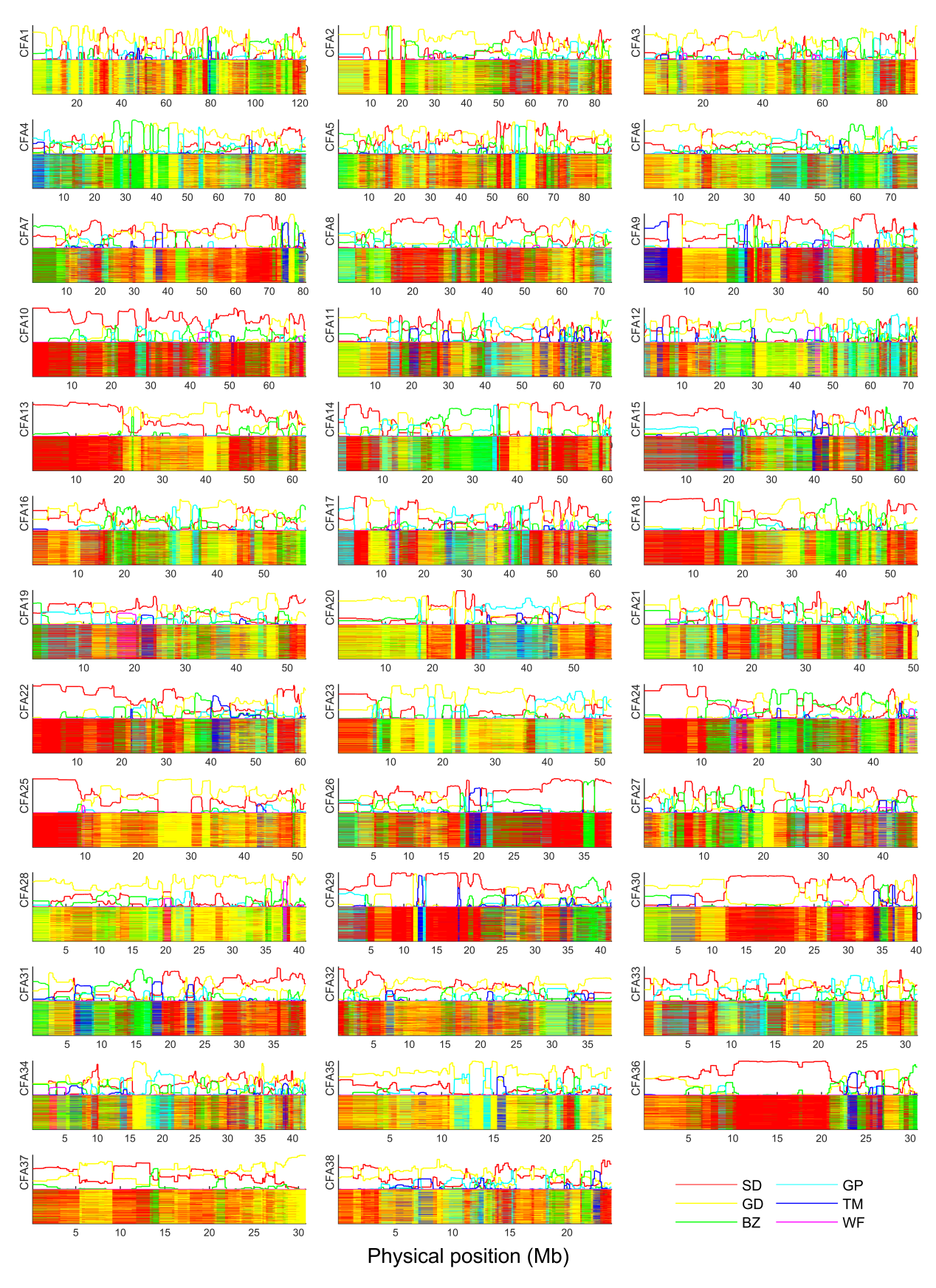

**Supplementary Fig. S1 | The inferred local ancestry along the genome of Irish Wolfhound.** In the line charts, the x-axis represents the physical positions on the chromosome, and the y-axis represents the local ancestral proportions. In the mosaic figures, the x-axis represents the physical positions on the chromosome, and the y-axis represents different haplotypes from the sample. Different colors represent different ancestors as shown in the legend. SD: Scottish Deerhound, GD: Great Dane, BZ: Borzoi, GP: Great Pyrenees, TM: Tibetan Mastiff, WF: Wolf.

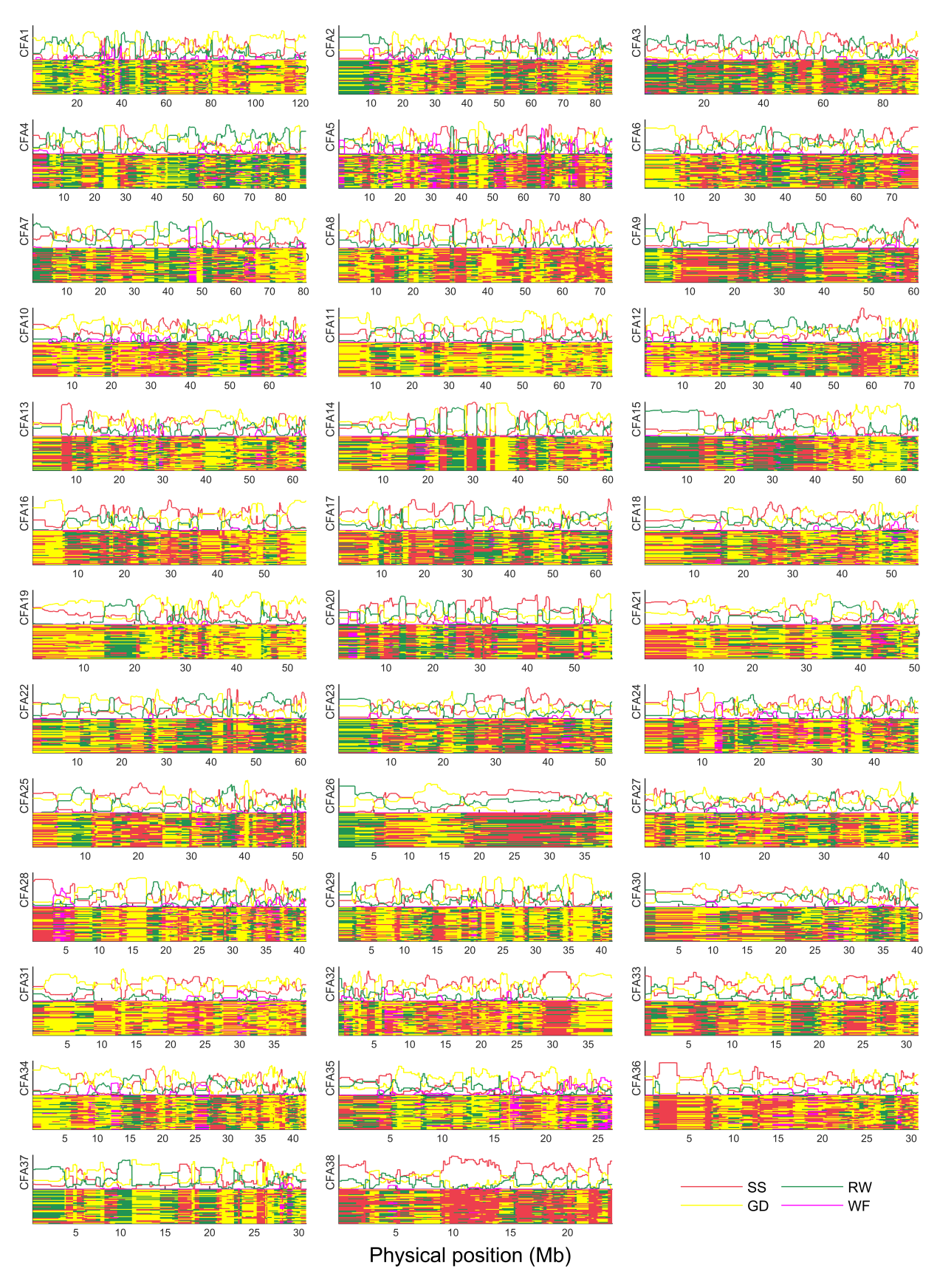

**Supplementary Fig. S2 | The inferred local ancestry along the genome of Giant Schnauzer.** In the line charts, the x-axis represents the physical positions on the chromosome, and the y-axis represents the local ancestral proportions. In the mosaic figures, the x-axis represents the physical positions on the chromosome, and the y-axis represents different haplotypes from the sample. Different colors represent different ancestors as shown in the legend. SS: Standard Schnauzer, GD: Great Dane, RW: Rottweiler, WF: Wolf.

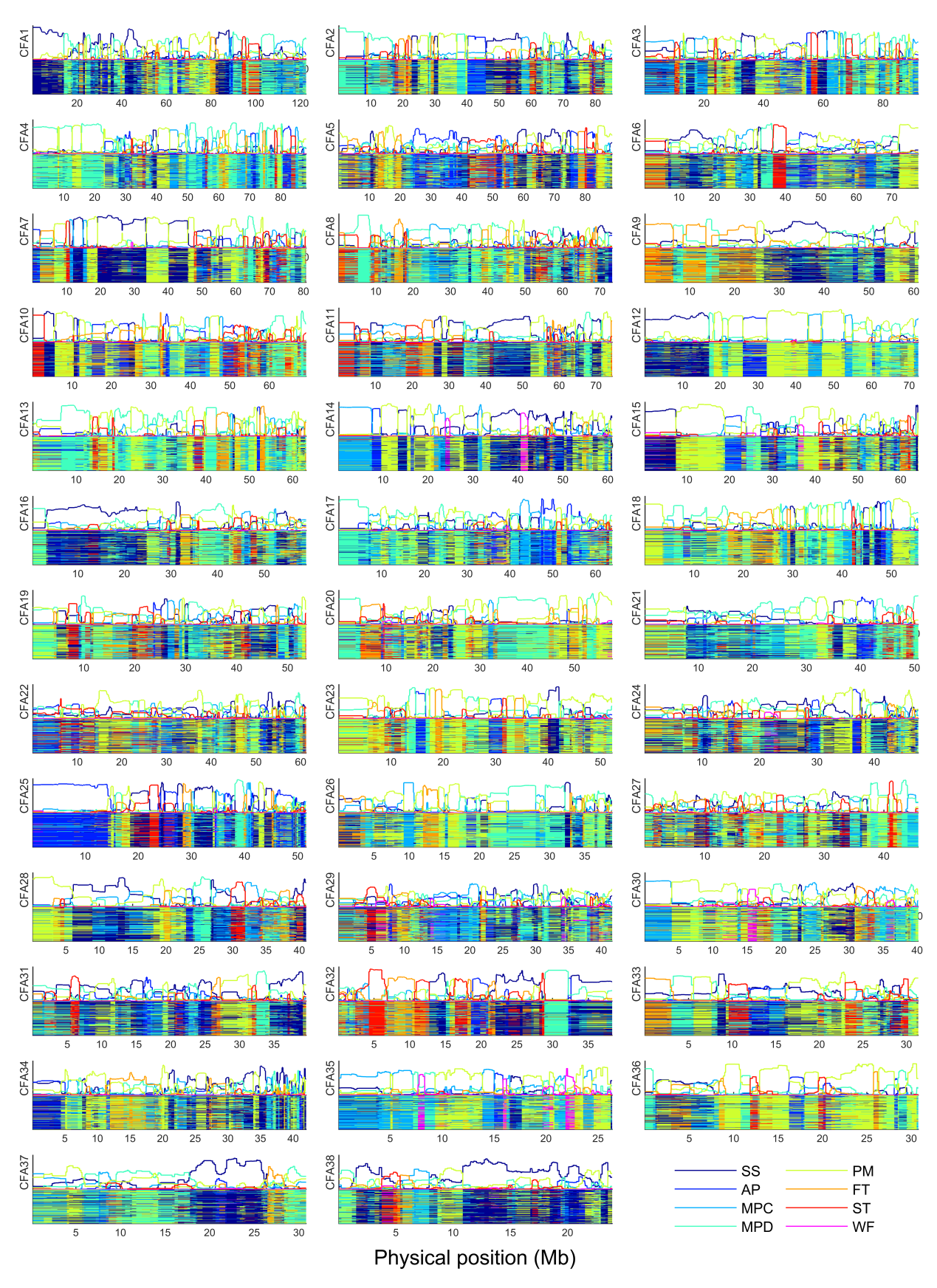

**Supplementary Fig. S3 | The inferred local ancestry along the genome of Miniature Schnauzer.** In the line charts, the x-axis represents the physical positions on the chromosome, and the y-axis represents the local ancestral proportions. In the mosaic figures, the x-axis represents the physical positions on the chromosome, and the y-axis represents different haplotypes from the sample. Different colors represent different ancestors as shown in the legend. SS: Standard Schnauzer, AP: Affenpinscher, MPC: Miniature Pinscher, MPD: Miniature Poodle, PM: Pomeranian, FT: Fox Terrier, ST: Scottish Terrier, WF: Wolf.

**2. Histograms of local ancestral proportions**

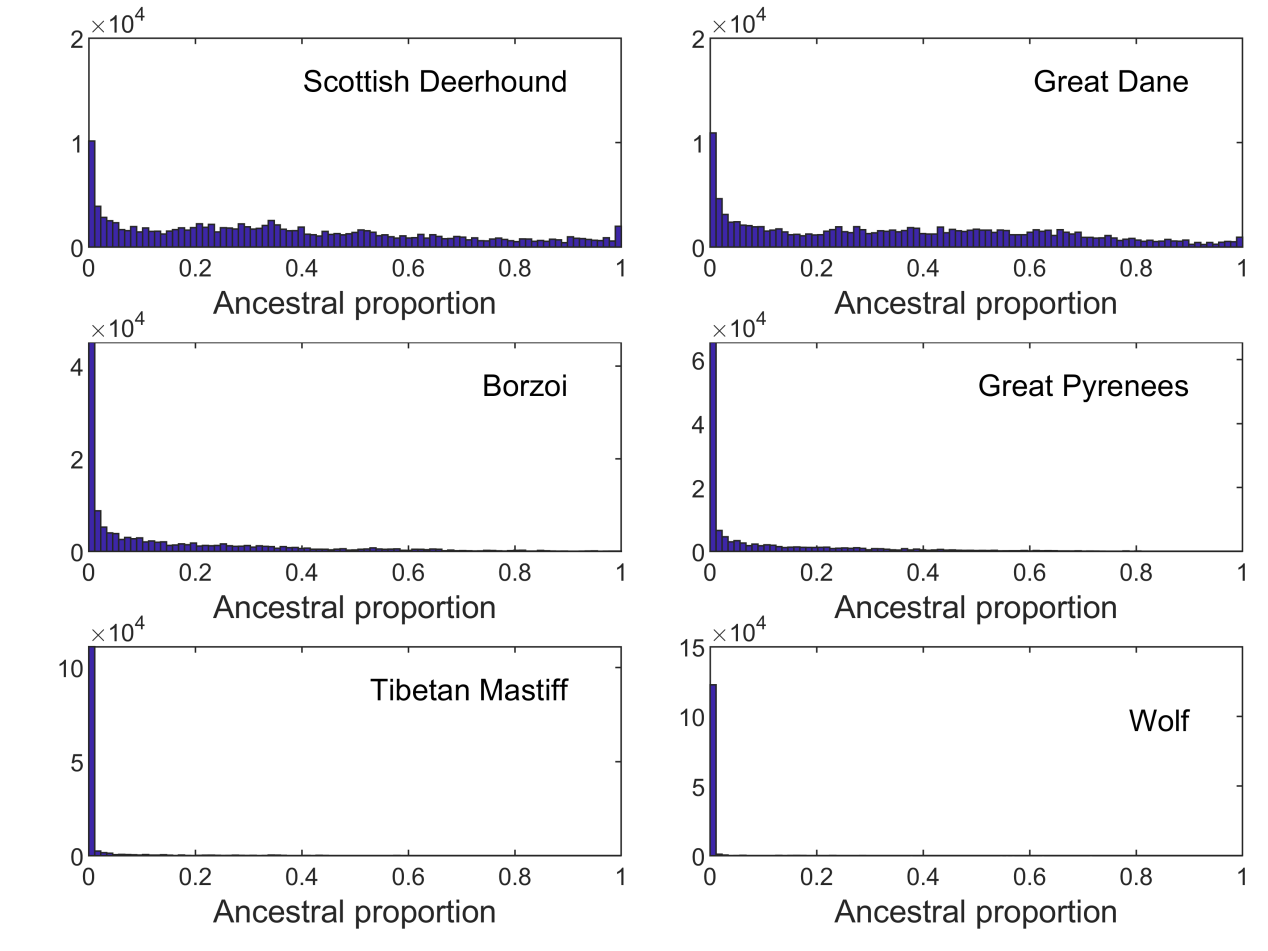

**Supplementary Fig. S4 | Histograms of local ancestral proportions of Irish Wolfhound**

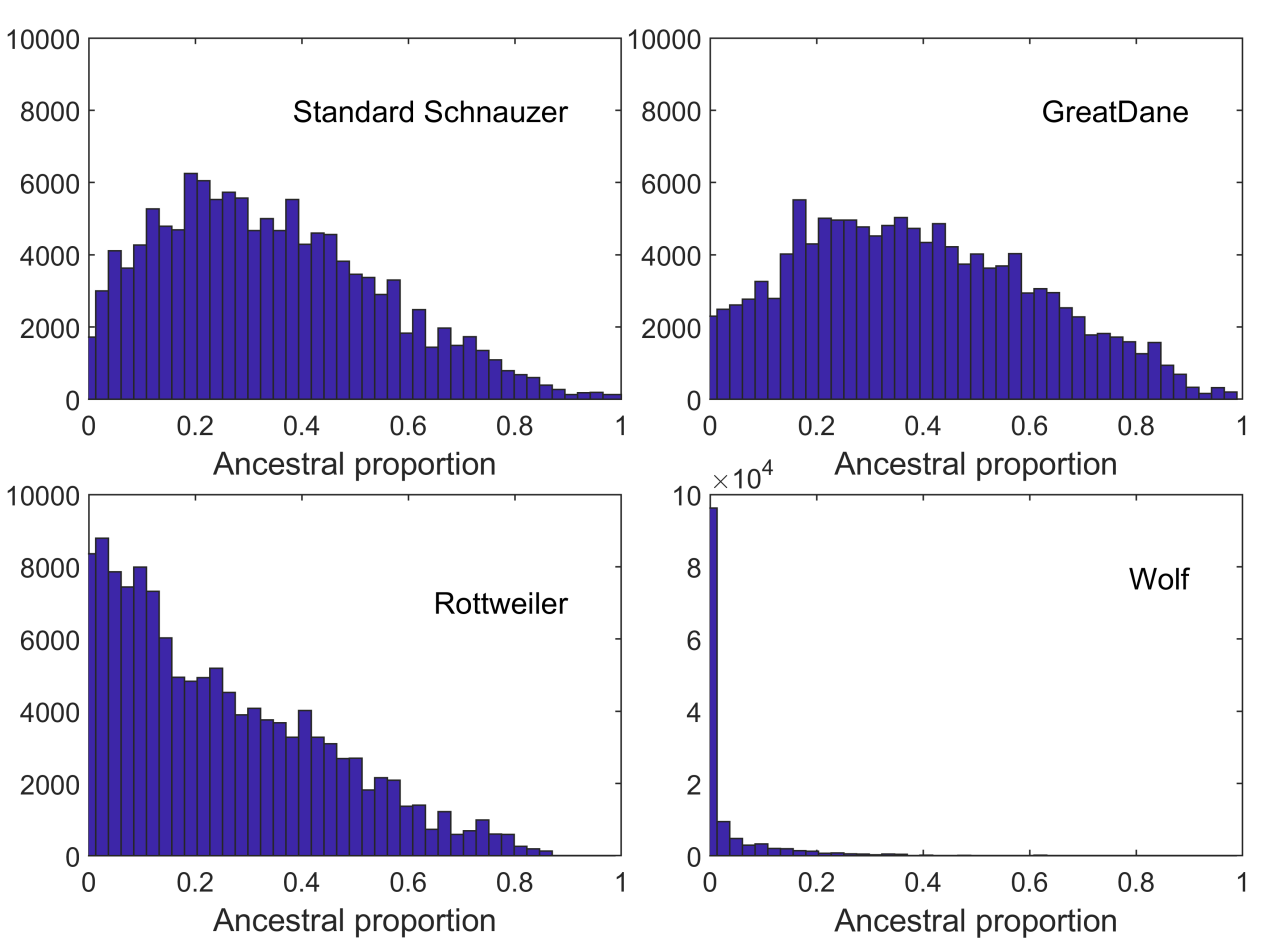

**Supplementary Fig. S5 | Histograms of local ancestral proportions of the Giant Schnauzer**

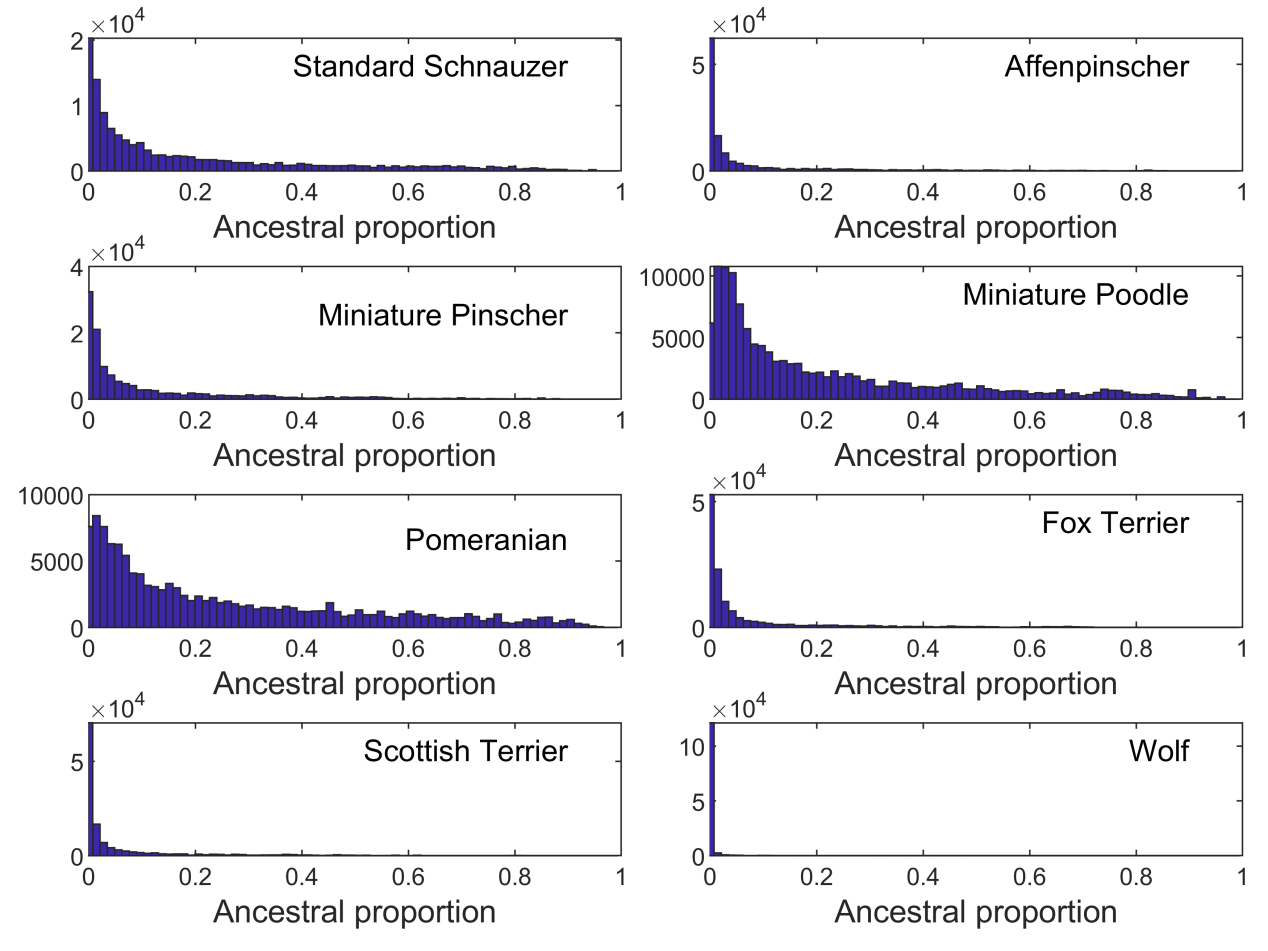

**Supplementary Fig. S6 | Histograms of local ancestral proportions of Miniature Schnauzer**

**3. Nucleotide diversity around the gene FGFR3**

A total of 1,629 dog samples representing ~500 dog breeds and 223 village dogs were used to calculate nucleotide diversity around the gene FGFR3 (Supplementary Fig. 7). The window size is chosen to be 2 kb and the step size 1 kb.

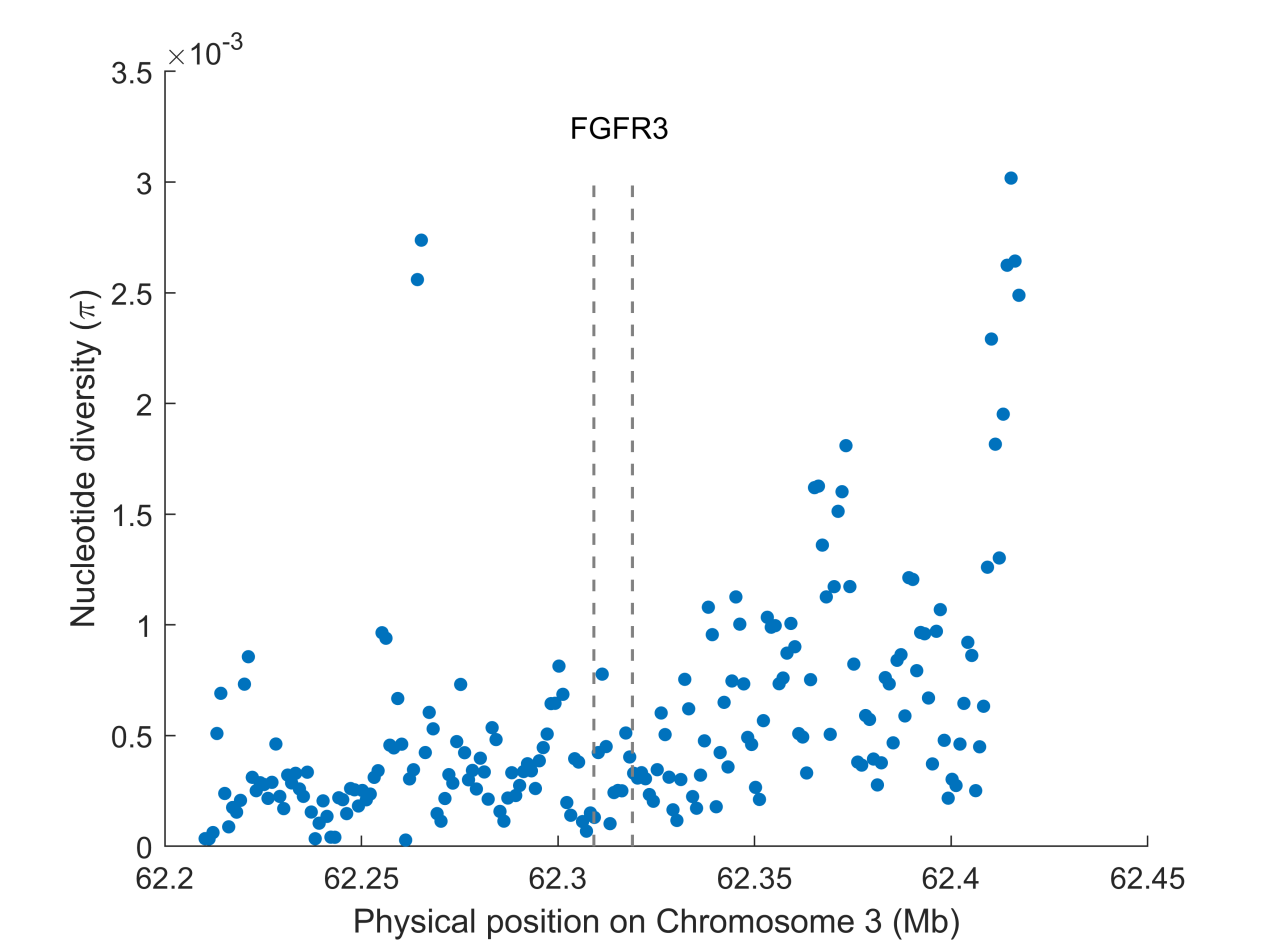

**Supplementary Fig. S7 | Nucleotide diversity around the gene FGFR3**

**4. Principal component analysis and outlier sample filtering**

We implemented SmartPCA from the EIGENSOFT package (version 6.1.4) to perform a principle component analysis. Loci with high linkage disequilibrium (r^2^>0.5) were filtered within a window with the size of 500kb and step size of 10. Outlier samples in the dashed ellipses were excluded from GAMA analysis, including 1 Scottish Deerhound (Supplementary Fig. 8), 2 Standard Schnauzer (Supplementary Figs. 9, 10), 4 Miniature Pinscher (Supplementary Fig. 10), and 1 Scottish Terrier (Supplementary Fig. 10).

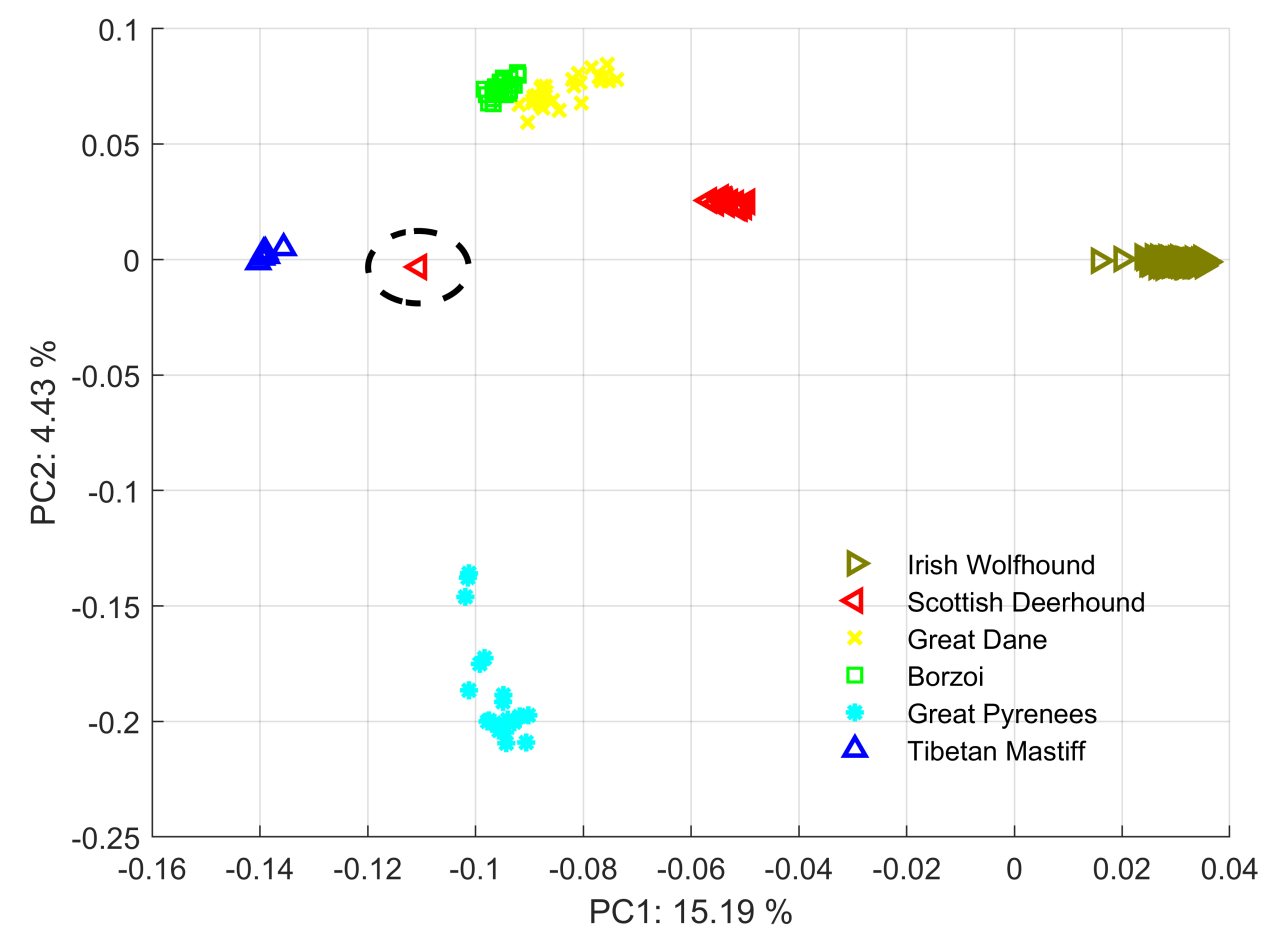

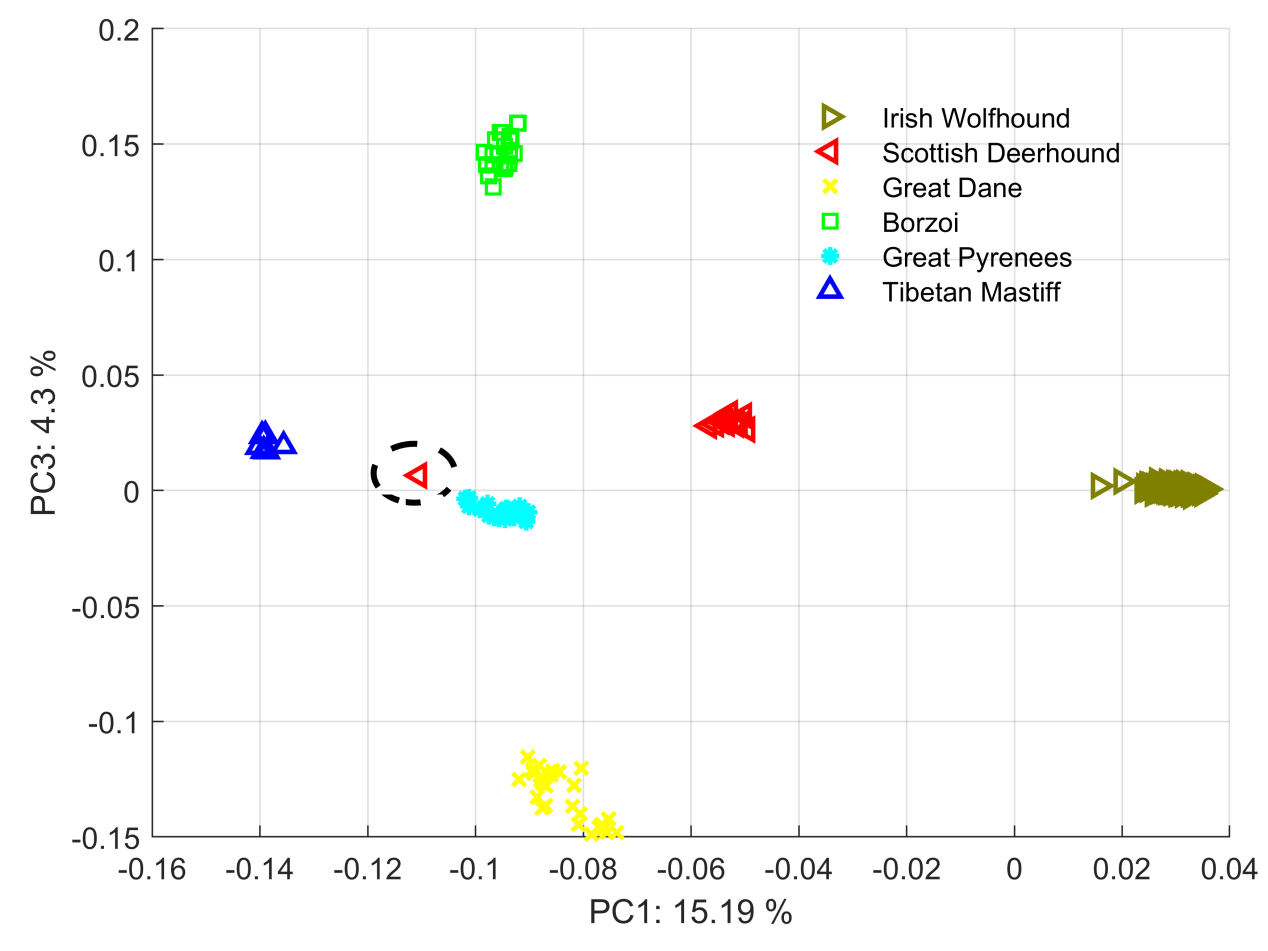

**Supplementary Fig. S8 | Principal component analysis of Irish Wolfhound and the five ancestral breeds.** SD: Scottish Deerhound, GD: Great Dane, BZ: Borzoi, GP: Great Pyrenees, TM: Tibetan Mastiff, IW: Irish Wolfhound. Outlier samples in the dashed ellipses are excluded from the GAMA analysis.

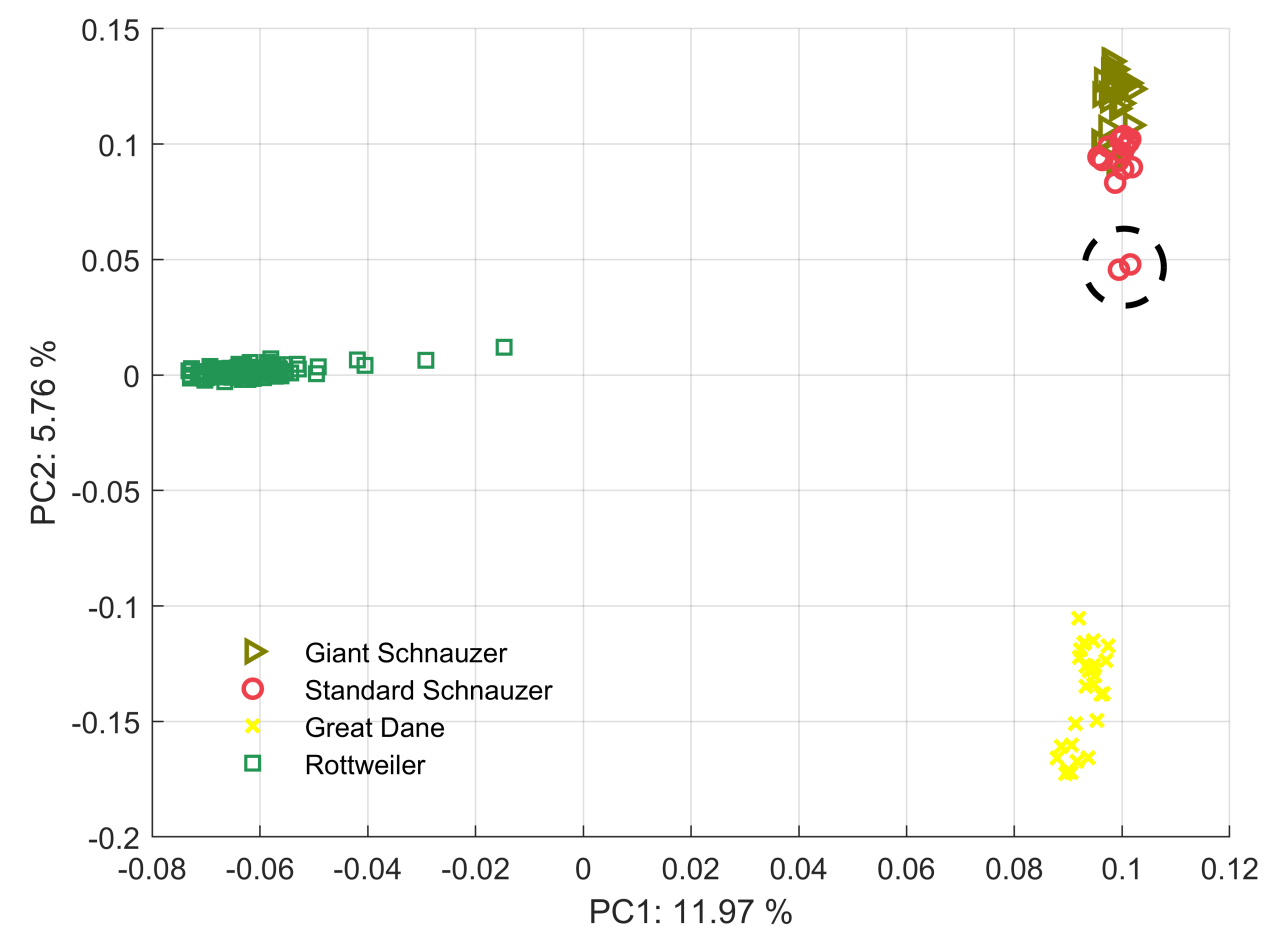

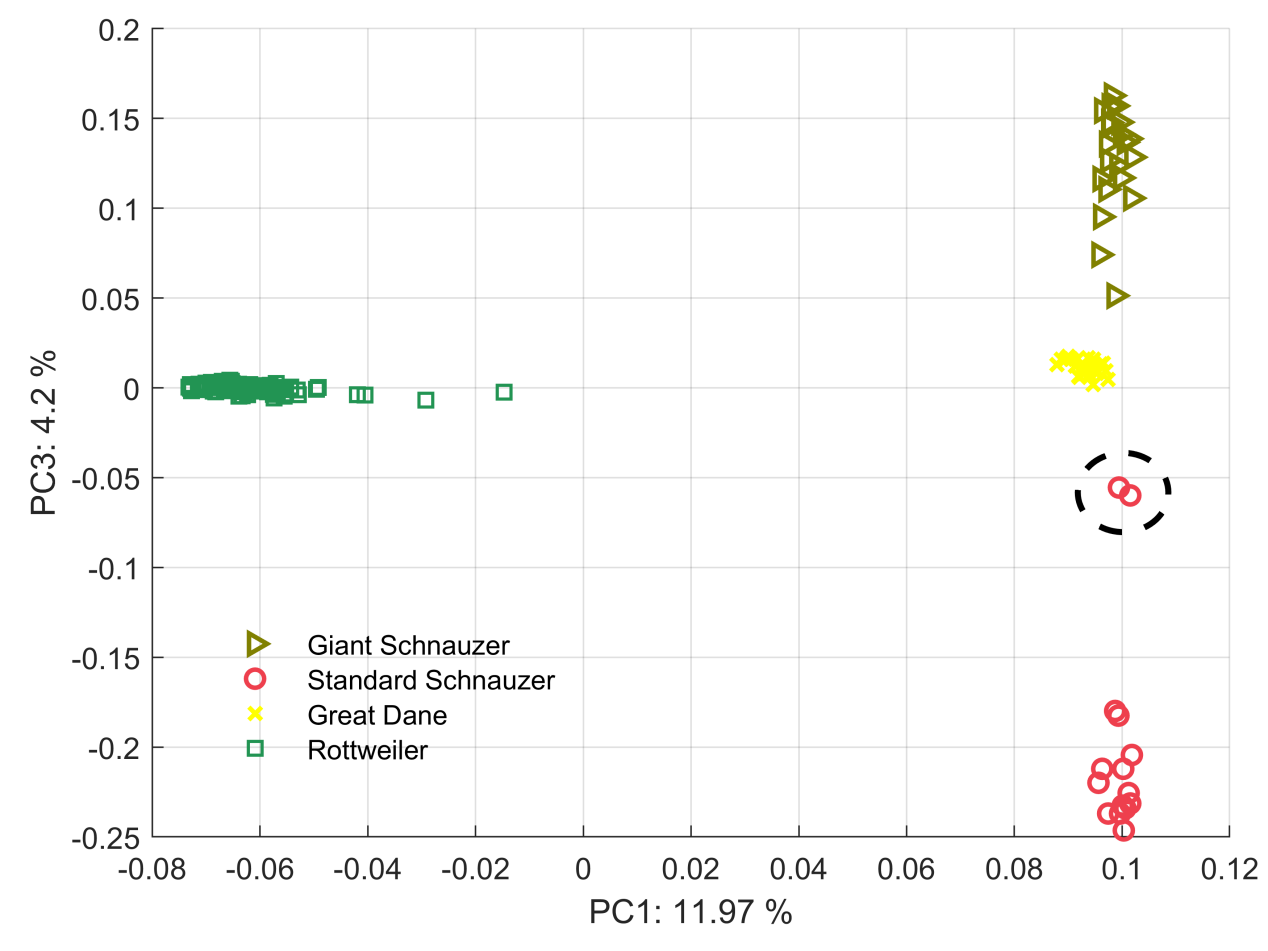

**Supplementary Fig. S9 | Principal component analysis of Giant Schnauzer and the three ancestral breeds.** SS: Giant Schnauzer, GD: Great Dane, RW: Rottweiler, GS: Giant Schnauzer. Outlier samples in the dashed ellipses are excluded from the GAMA analysis.

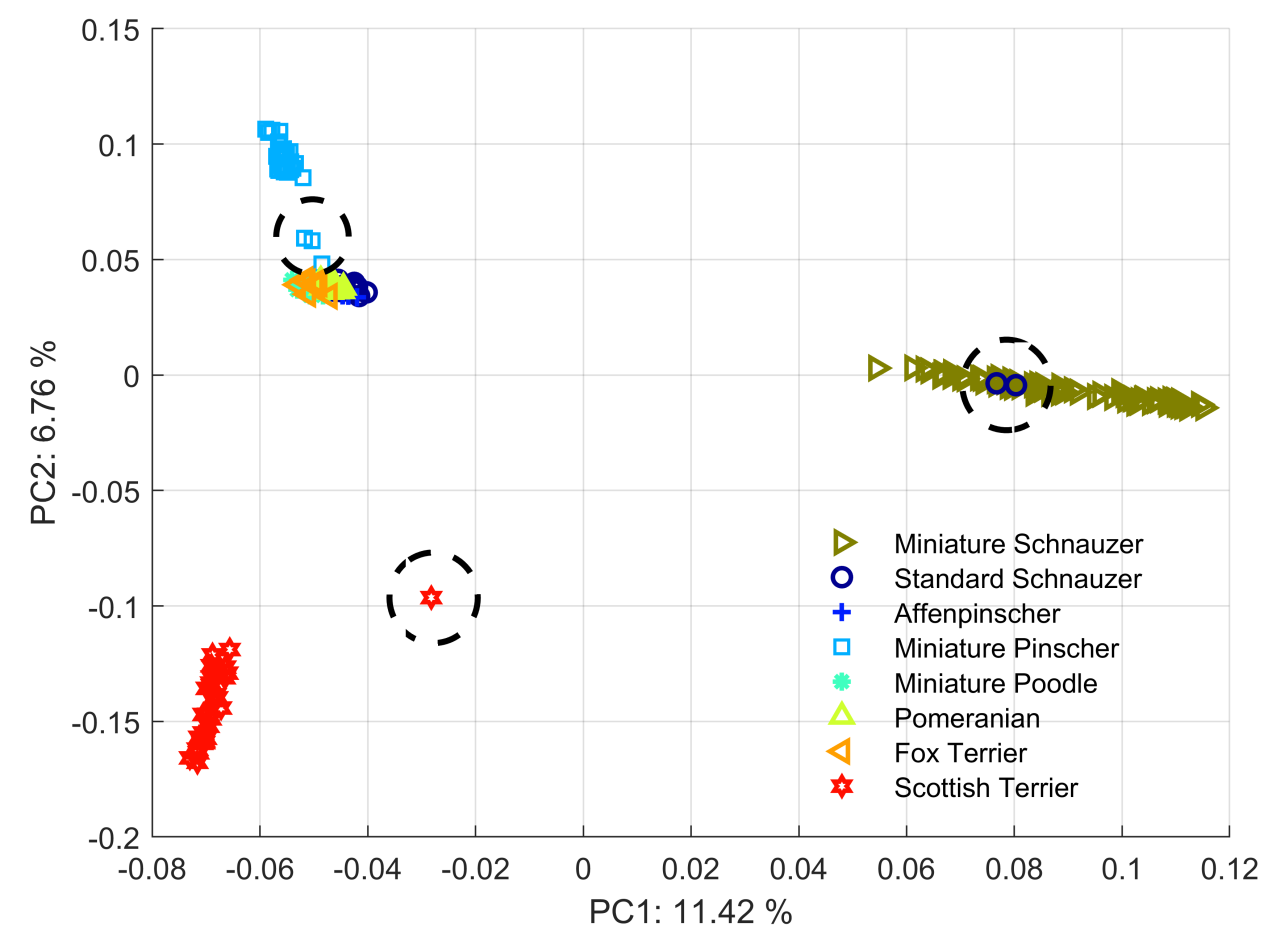

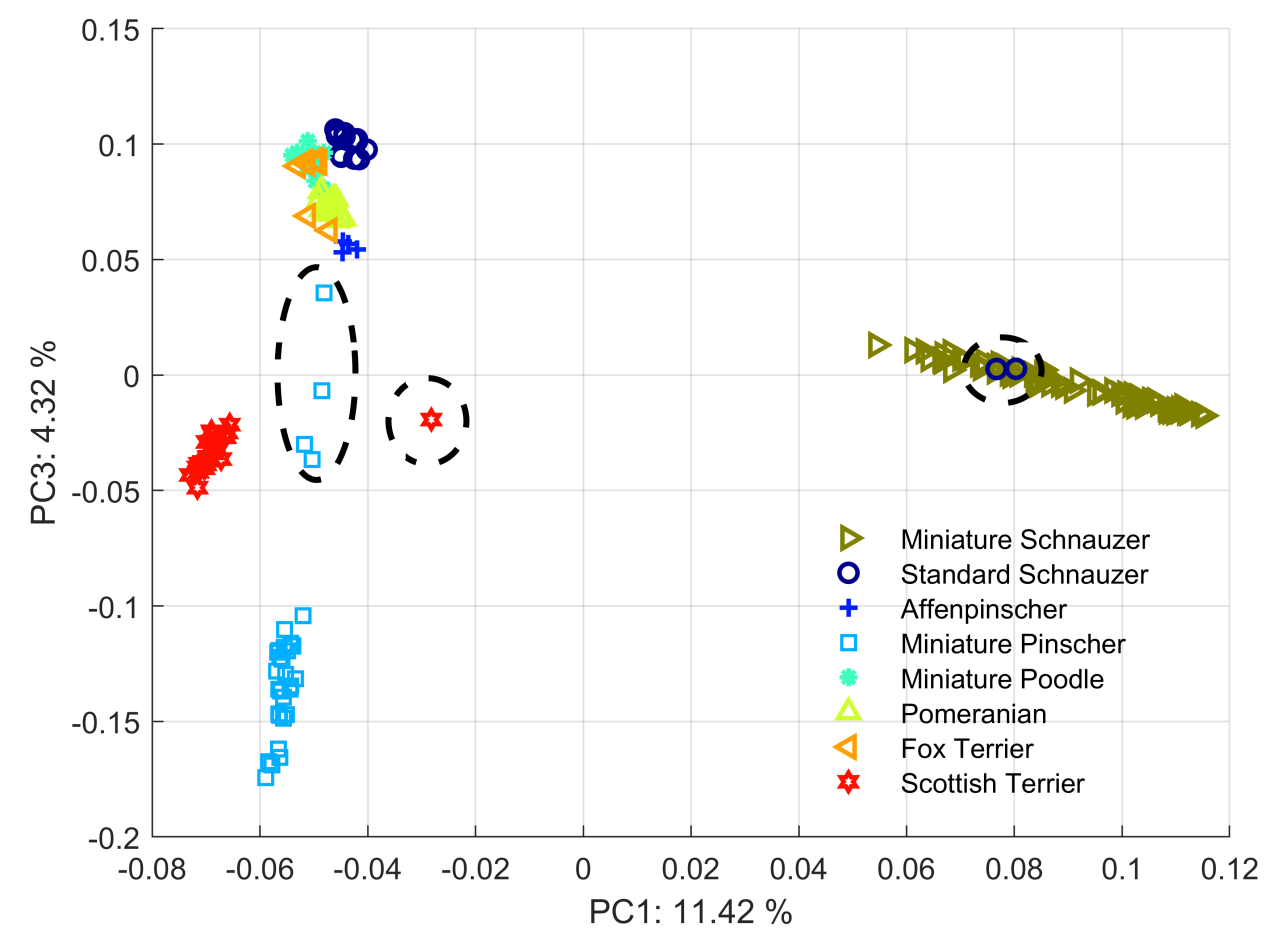

**Supplementary Fig. S10 | Principal component analysis of Miniature Schnauzer and the seven ancestral breeds.** SS: Standard Schnauzer, AP: Affenpinscher, MPC: Miniature Pinscher, MPD:	Miniature Poodle, PM: Pomeranian, FT: Fox Terrier, ST: Scottish Terrier, MS: Miniature Schnauzer. Outlier samples in the dashed ellipses are excluded from the GAMA analysis.

**5. Candidate artificial selection regions with multiple genes**

*CTGF* and *FAM210A* in purified segments of chromosome 1 exhibit molecular functions relevant to body size. Ctgf plays an essential role in chondrocyte proliferation, differentiation, and cell adhesion in many cell types. It is widely expressed during development, and in adulthood, it is expressed in all major organs, including the skeletal system ^1, 2^. Ctgf ^Hi/+^ animals had significantly smaller bone area and lean mass than controls exhibiting a smaller overall body size. Ctgf ^Hi/+^ animals had low bone mineral density, suggesting changes in bone mineralization or morphology ^3^. *FAM210A* was demonstrated to have a crucial role in regulating the bone structure and function ^4^.

*WAC* and *BAMBI* in purified segments of chromosome 2 all exhibit molecular functions relevant to body size. *WAC* loss-of-function mutations cause DeSanto-Shinawi syndrome, characterized by global developmental delay that is apparent in infancy or early childhood ^5^. *BAMBI* seems to be co-expressed with *BMP4* and supposedly acts as a negative feedback loop to control BMP activity ^6^. BMPs participate in a negative feedback loop with FGFs in the developing limb. The inhibition of FGFs by BMPs is significant for activating Sox9 expression, which is essential for chondrogenesis of the developing skeletal tissue in avian and mammalian limb buds ^7^. The negative feedback between BMP and FGF signaling is also present in the growth pates of mammalian long bones ^7^. Bambi ^-/-^ mouse had a decreased weight in females ^8^.

Five genes involved in purified segments of chromosome 4 show evidence relevant to body size. *STC2*, a secreted glycoprotein hormone, inhibits growth in mice independently of the GH/IGF1 pathway ^9, 10^. *STC2* plays a role in calcium and phosphate homeostasis ^11^. *STC2* knock-out mice are larger and grow faster than wild-type controls ^10^. Several studies have reported the correlation of *STC2* with domestic dog body size ^12, 13^. Homeobox transcription factor Nkx2-5, highly expressed in the heart, is critical during early embryonic cardiac development. A missense mutation in the transcription factor *NKX2-5* can cause hypothyroidism and growth delay in humans ^14^. Nkx2-5 mutant mice exhibit phenotypes of embryonic growth retardation and decreased embryo size ^15, 16, 17^. *CREBRF* significantly correlates with human body height ^18^. A missense variant in *CREBRF* strongly influences body mass index in Samoans ^19^. The *CREBRF* knock-out mice exhibit decreased body weight ^20^. *DUSP1*, a nucleus-localized MKP, is a central negative regulator of the MAPK pathway and participates in maintaining homeostasis of glucose metabolism and energy balance in peripheral tissues. Its activation facilitates obesity by lowering energy expenditure. *DUSP1* deficient protected mice from diet-induced obesity ^21^.

Eleven genes involved in purified segments of chromosome 6 show evidence relevant to body size. HspB1^-/-^ mice had delayed growth with significantly lower body weight than controls ^22^. Some authors postulate that reduced expression of genes encoding intracellular HSPs, mainly *HSPA1A* and *HSPB1*, may contribute to metabolic syndromes such as obesity and diabetes ^23, 24^. High levels of *HSPB1* mRNA are associated with enhanced insulin sensitivity in skeletal muscle ^25^. Decreases in *HSPB1* mRNA and intracellular protein can affect adipose tissue and could be associated with fat reduction, and a decrease in expression of *HSPB1* may influence glucose sensitivity ^26^. *ZP3* is a differentially expressed gene associated with differential body size in mandarin fish ^27^. Several studies in rodents have given evidence for the role of *VGF* in controlling energy metabolism ^28, 29^ and food intake ^30^. Targeted deletion of *VGF* produces a lean, small, and hypermetabolic mouse ^29^. *POR*-dependent cholesterol synthesis is essential during limb and skeletal development ^31^. Some patients with *POR* mutations have defects in the limb skeleton ^32, 33^. Studies also revealed that the genes *CUX1* ^34, 35, 36^, *MDH2* ^37^, *PLOD3* ^38^, *SERPINE1* ^39, 40^, *SRRM3* ^41^, *TRIM56* ^37^, and *ZNHIT1* ^42^ are implicated in developmental regulation, exhibiting various phenotypes such as growth retardation, delayed limb development, short tail, abnormal body weight, and decreased body length.
